# Conserved cellular architecture and developmental mechanisms of the zebrafish meninges

**DOI:** 10.64898/2026.02.14.703877

**Authors:** Ashley L. Arancio, Kathryn Wilhem, Hung-Jhen Chen, Brandon M. Hernandez, Percy J. Raggi, D’Juan T. Farmer

## Abstract

The meninges are a multilayered connective tissue that supports and protects the brain and skull, yet their developmental origins and signaling functions remain poorly understood. Here, we establish zebrafish as a tractable model for defining meningeal development and function across larval and adult stages. Using a meninges-enriched *foxc1b*:Gal4 reporter, we resolve the spatiotemporal emergence of meningeal fibroblasts and demonstrate a conserved, dosage-sensitive requirement for Foxc1 activity during meningeal formation. A targeted pharmacological screen identifies Wnt signaling as essential for timely establishment of the primary meninx. To define adult meningeal diversity, we integrate single-cell multiome profiling with spatial validation and identify multiple transcriptionally distinct populations organized into layered compartments, including pial, arachnoid, dural, and periosteal dura fibroblasts. Finally, inducible larval ablation of meningeal cells reveals limited regenerative capacity following widespread loss and leads to persistent defects in calvarial osteogenesis and brain architecture, including reduced osteoblast differentiation at bone fronts and disrupted tissue organization at sites lacking meningeal recovery. Together, these findings define key features of zebrafish meninges and provide a framework for dissecting meningeal development, regeneration, and meninges-dependent signaling in vivo.

**Summary Statement:** This study uses zebrafish to show how the tissues surrounding the brain develop early and guide proper formation of both the brain and skull.

## Introduction

The meninges are a multilayered connective tissue that encase the central nervous system and, in the head, lie between the brain and the skull. They are comprised of transcriptionally distinct fibroblast populations, with resident immune cells and an extensive vasculature that support tissue nourishment and waste removal (Dasgupta and Jeong, 2019; Decimo et al., 2012; O’Rahilly and Muller, 1986). Meningeal fibroblasts are broadly organized into three layers: the pia mater, which lies adjacent to the brain and is separated from neural tissue by a basement membrane; the arachnoid mater, which contributes to cerebrospinal fluid regulation; and the dura mater, which anchors the meninges to the overlying skull and serves as a critical immunological niche.

Recent single-cell sequencing studies have begun to resolve the cellular composition of embryonic and adult mouse and human meninges (Degenhard et al., 2024; DeSisto et al., 2020; Ebert et al., 2025; Farmer et al., 2021; Kearns et al., 2023; Li et al., 2026; Pietila et al., 2023; Sun et al., 2025; Vinsland et al., 2025). These analyses reveal substantial fibroblast heterogeneity by embryonic day 14 (E14) in mice and gestational week 8 in humans and extensive diversity within both the leptomeninges (pia and arachnoid) and the dura mater.

Beyond their structural role, the meninges regulate the development of neighboring tissues, including the brain and the skull (Como et al., 2023; Dasgupta and Jeong, 2019). Meningeal-derived retinoic acid signaling modulates cortical neurogenesis (Como et al., 2025; Haushalter et al., 2017; Matrongolo et al., 2023; Mishra et al., 2016; Siegenthaler et al., 2009), while TGFβ family ligands secreted from meninges (Greenwald et al., 2000; Slater et al., 2009) influence cranial suture maintenance and calvarial growth (Cooper et al., 2012; Greenwald et al., 2000; Kwan et al., 2008; Opperman et al., 1993; Slater et al., 2008). Despite these roles, the mechanisms governing meningeal development remain incompletely understood. The forkhead box transcription factor FOXC1 is required for meningeal formation in mice and humans, as hypomorphic or null alleles disrupt meningeal development (Aldinger et al., 2009; Siegenthaler et al., 2009; Vivatbutsiri et al., 2008; Zarbalis et al., 2007). Similarly, ZIC family transcription factors are required for meningeal development, as Zic1/3 double mutants exhibit basement membrane defects and impaired fibroblast proliferation (Inoue et al., 2008). In both contexts, genetic perturbations broadly affect all meningeal layers, and insights into cell-type specific developmental programs are just emerging (Derk et al., 2023).

Although most functional studies of the meninges have used rodents, ultrastructural analyses reveal stratified meningeal layers across vertebrate and invertebrate species, suggesting deep evolutionary conservation (Kappers, 1925; Nakao, 1979). Zebrafish also possess stratified meninges containing vasculature and immune cell populations comparable to those of mammals (Galanternik et al., 2025). However, the relationships between zebrafish and mammalian meningeal layers, the transcriptional diversity of zebrafish meningeal fibroblasts, and the conservation of meningeal signaling functions remain unresolved.

Here, we define the developmental timing and molecular identities of meningeal fibroblasts in zebrafish. Using *foxc1a* and *foxc1b* mutant alleles, we demonstrate a conserved, dosage-sensitive requirement for FOXC1 during meningeal formation. Pharmacological perturbation identifies Wnt signaling as essential for timely establishment of the embryonic meninx, while single-cell genomics resolves transcriptionally distinct meningeal layers with parallels to murine and human populations. Finally, larval meningeal ablation reveals limited regenerative capacity following large-scale loss and persistent defects in brain and skull development. Together, these findings establish zebrafish as a powerful model for dissecting developmental and regulatory programs governing meningeal biology.

## Results

### A *foxc1b*:Gal4 reporter marks developing meningeal populations in zebrafish

To assess whether the zebrafish paralogs for Foxc1 (*foxc1a* and *foxc1b*) are active in meninges, we examined zebrafish carrying the *foxc1b*:Gal4;UAS:mCherryNTR transgene (Whitesell et al., 2019). In contrast to the broadly expressed *foxc1b*:GFP reporter, mCherry fluorescence was enriched around the brain (**Fig. 1A**). Both reporters are driven by the same promoter, suggesting that their differences reflect different genomic integration sites. Orthogonal sections revealed co-detection of mCherry and GFP signals within a single cellular layer adjacent to the brain (**Fig. 1A, Table S1**). Imaging with the epithelial marker *cdh1*:mlanYFP confirmed that mCherry⁺ cells reside beneath the epithelium (**Fig. 1B, Table S1**). To assess mesenchymal identity, *foxc1b*:Gal4;UAS:mCherryNTR fish were crossed with *fli1a*:GFP, a broad marker of craniofacial mesenchyme. mCherry⁺ cells encasing the brain co-expressed *fli1a*:GFP, whereas *fli1a*:GFP-labeled vasculature lacked detectable mCherry signal (**Fig. 1C, Table S1**). A developmental time course revealed no detectable mCherry signal at 24 hpf. By 36–48 hpf, mCherry⁺/*fli1a*:GFP⁺ cells emerged within the rostral meninges, while *fli1a*:GFP⁺ cells preceded mCherry detection caudally (**Fig. 1D**). Time-lapse imaging confirmed early mCherry activation near the rostral hindbrain, followed by expansion toward the midbrain. Ventral populations expanded toward both the dorsal-lateral and dorsal-caudal regions, with *fli1a* expression preceding mCherry in all regions (**Movie 1**). Transient mCherry expression was also detected in muscle and yolk-associated cells that was not observed at later developmental stages. Together, these data define the spatiotemporal emergence of *foxc1b*⁺ meningeal cells and establish *foxc1b*:Gal4 as a meninges-enriched reporter.

**Figure 1.**
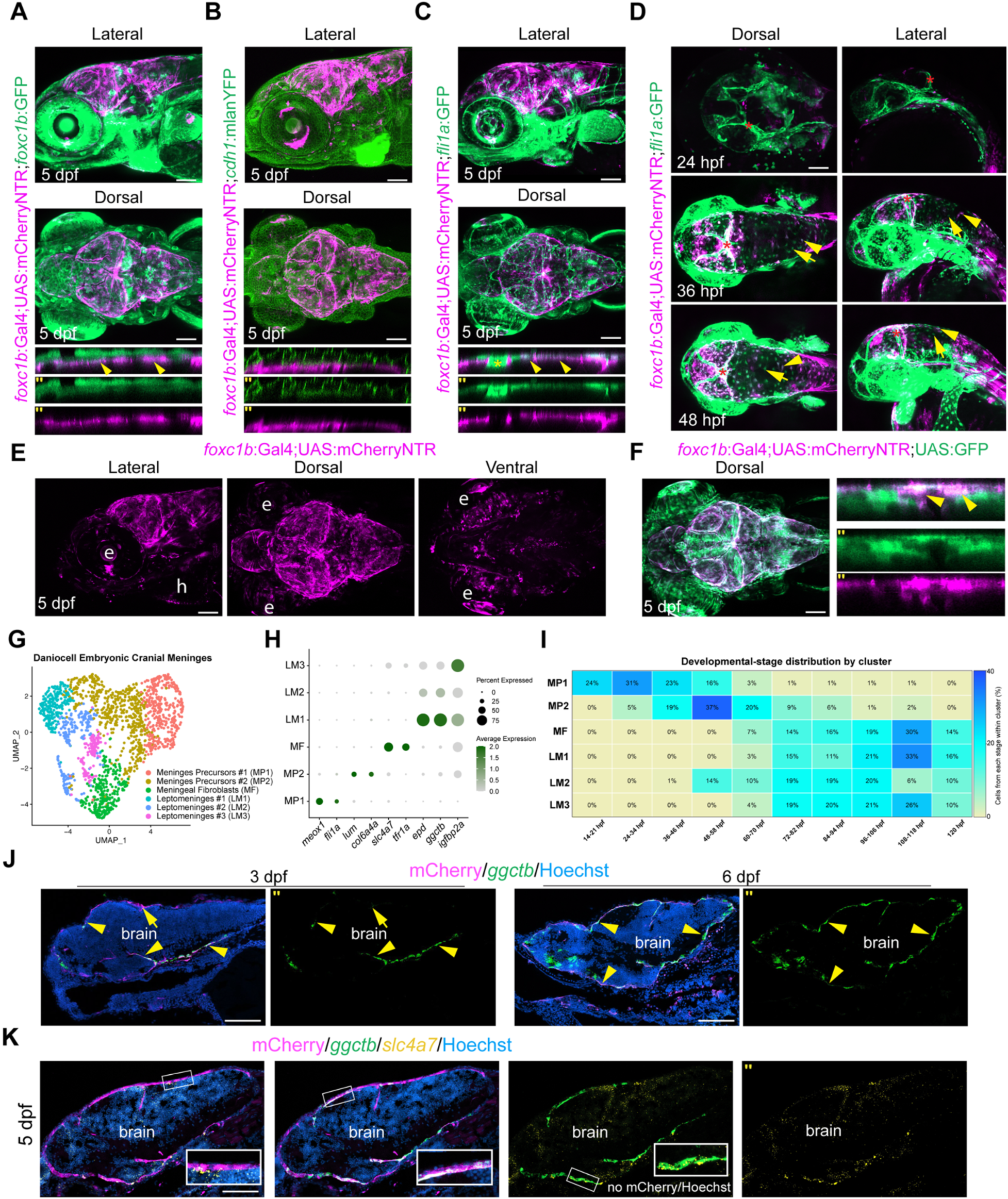
A *foxc1b* reporter labels larval meninges and reveals early spatial patterns of meningeal formation. **(A–C)** Confocal images of 5 dpf *foxc1b*:Gal4;UAS:mCherryNTR larvae with additional transgenic markers. Lateral (top) and dorsal (bottom) views and orthogonal sections are shown (n = 5 per condition). **(A)** *foxc1b*:GFP. Arrowheads indicate GFP⁺mCherry⁺ cells surrounding the brain. **(B)** Epithelial marker *cdh1*:mlanYFP. **(C)** Mesenchymal marker *fli1a*:GFP. Arrowheads indicate double-positive cells. Asterisk marks blood vessel. **(D)** Developmental time course (24–48 hpf) of *fli1a*:GFP and mCherry expression. Arrowheads denote mCherry⁺GFP⁺ cells and arrows indicate GFP^+^-only cells (n = 5 per stage). Asterisks mark representative blood vessels. **(E)** Lateral, dorsal, and ventral views of *foxc1b*:Gal4;UAS:mCherryNTR larvae (n = 11). h = heart, e = eye. **(F)** Dorsal view and orthogonal section of *foxc1b*:Gal4;UAS:mCherryNTR; UAS:GFP larvae (n = 5). Arrowheads indicate mCherry⁺GFP⁺ meningeal cells. **(G)** UMAP of subsetted meningeal populations from the Daniocell single-cell RNA-sequencing dataset (Sur et al., 2023). **(H)** Dot plot of marker genes enriched in each meningeal cluster. **(I)** Heatmap showing cluster composition by developmental stage. **(J)** RNAscope for the leptomeningeal marker *ggctb* with mCherry immunostaining at 3 and 6 dpf (n = 3). Arrowheads denote mCherry⁺*ggctb*⁺ cells. **(K)** RNAscope for *ggctb* and the meningeal fibroblast marker *slc4a7* with mCherry immunostaining (n = 3). Insets show higher-magnification views highlighting co-detection. (’’) = single channels. Scale bars, 100 µm.

To define the broader distribution of mCherry⁺ cells, we performed whole-body and head-specific imaging of *foxc1b*:Gal4;UAS:mCherryNTR fish at 5 dpf and observed scattered expression in the heart, fins, eyes, lower jaw and trunk (**Fig. 1E**, **Fig. S1A-C**), consistent with previous reports using this reporter to visualize other cell types, such as vascular smooth muscle cells (Whitesell et al., 2019). Comparison of UAS:mCherryNTR and UAS:GFP reporters revealed broad GFP expression throughout the body, including the meninges, in contrast to meningeal enrichment of mCherry (**Fig. 1F**, **Fig. S1A-C, Table S1**). Thus, while all lines showed meningeal expression, the combination of *foxc1b*:Gal4 and UAS:mCherryNTR preferentially enriches transgene expression within the meninges, a feature exploited throughout this study.

Next, to resolve the transcriptional composition of embryonic meninges, we subsetted and reclustered previously annotated meningeal populations from the Daniocell single-cell transcriptomic dataset (Sur et al., 2023). This resolved two clusters of meningeal precursors, one cluster of meningeal fibroblasts, and three clusters of leptomeninges (**Fig. 1G**). Precursors expressed mesenchymal markers including *fli1a, lum*, and *meox1*, while leptomeningeal clusters expressed *epd* and *ggctb* and meningeal fibroblasts were distinguished by *slc4a7* and *tfr1a* (**Fig. 1H**) (Galanternik et al., 2025). Precursor clusters were enriched for cells from 14–70 hpf, whereas fibroblast and leptomeningeal clusters were primarily composed of cells from 72 hpf and later, consistent with developmental transitions between these states (**Fig. 1I**). Consistent with progressive differentiation, *ggctb* expression was primarily ventral at 3 dpf but expanded throughout the meninges by 6 dpf (**Fig. 1J**). Co-staining with *slc4a7* and *ggctb* at 5 dpf to differentiate fibroblast and leptomeningeal cell types revealed single- and double-positive mCherry⁺ meningeal cells that did not form discrete layers (**Fig. 1K, Table S1**). These findings support a heterogeneous but unstratified meninges at larval stages.

### Foxc1 paralogs exhibit conserved requirements during meningeal development

To test conserved requirements for meningeal development, we examined *foxc1a* and *foxc1b*, which were expressed across meningeal precursor, fibroblast, and leptomeningeal populations (**Fig. 2A**). RNAscope in situ hybridization at 24 hpf revealed few dorsal *foxc1a*⁺ or *foxc1b*⁺ cells and minimal mCherry around the developing brain, with most *foxc1* transcripts enriched ventrally near the pharyngeal arches (**Fig. 2B**). By 36 hpf, mCherry⁺ cells covered the ventral brain surface and were also present dorsally near the midbrain-hindbrain boundary (**Fig. 2B, Table S1**). Both *foxc1* paralogs extended beyond the mCherry domain toward the forebrain, suggesting that reporter accumulation follows transcriptional activation. By 5 dpf, mCherry and both paralogs were broadly co-expressed throughout the meninges, although a small population of mCherry-negative, *foxc1a*/*foxc1b*-positive cells were detected around the forebrain (**Fig. 2B, Table S1**). In some regions surrounding the brain, *foxc1a* was detected in the absence of *foxc1b*. At both stages, *foxc1a*/*b* expression was detected outside the meninges, including throughout the pharyngeal arches and lower jaw, where it did not overlap with mCherry.

**Figure 2.**
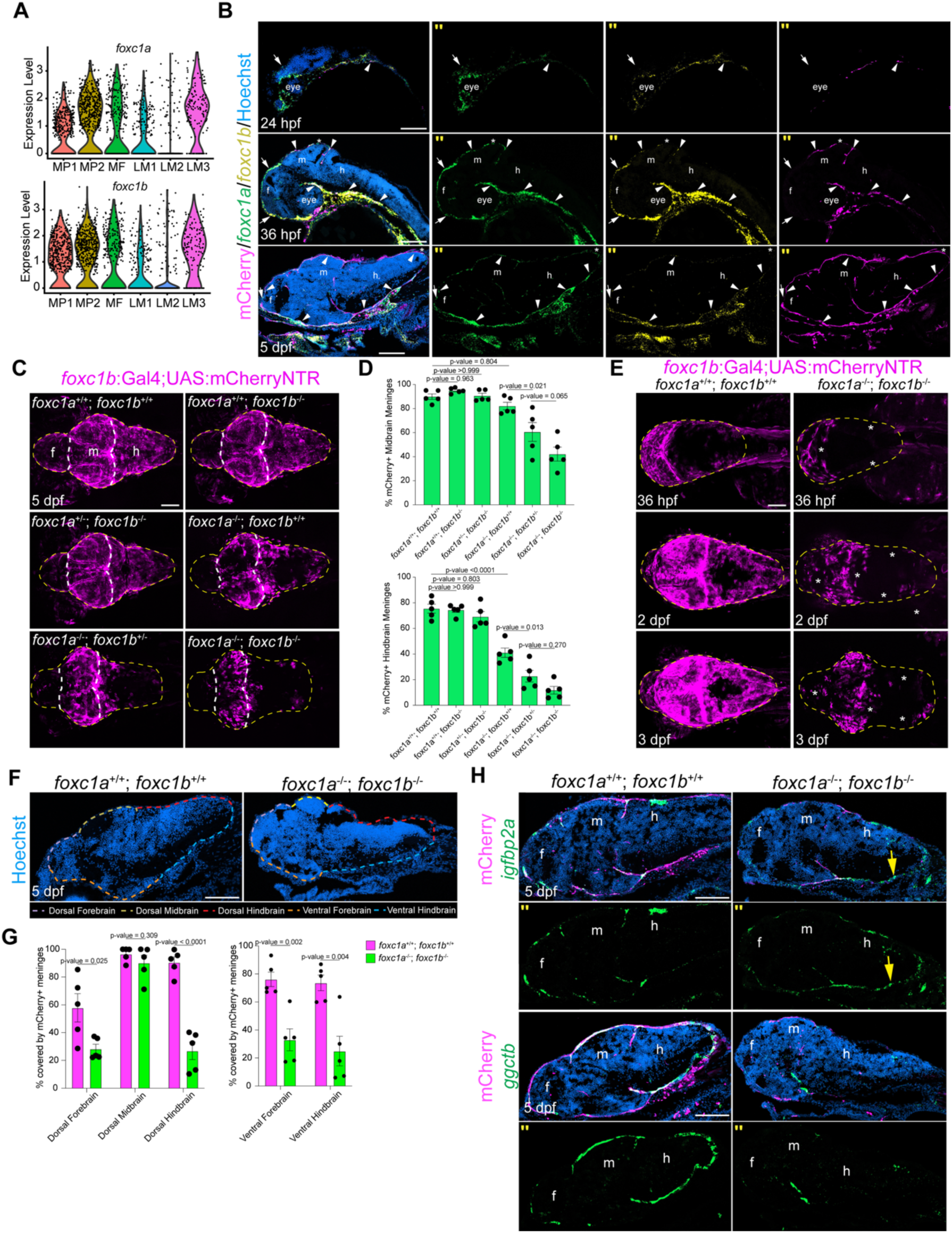
*foxc1a* and *foxc1b* exhibit dosage-sensitive requirements for meningeal formation. **(A)** Violin plots of *foxc1a* and *foxc1b* expression across meningeal clusters from Fig. 1G. **(B)** RNAscope for *foxc1a* and *foxc1b* with mCherry immunostaining in *foxc1b*:Gal4;UAS:mCherryNTR fish at 24 hpf, 36 hpf, and 5 dpf (n = 3 per stage). Arrowheads indicate *foxc1a*^+^*foxc1b*^+^mCherry^+^ cells and arrows indicate *foxc1a*^+^*foxc1b*^+^mCherry^-^ cells. Asterisks indicate *foxc1a*^+^*foxc1b*^-^ cells. f = forebrain, m = midbrain, h = hindbrain. **(C)** Dorsal confocal images of *foxc1b*:Gal4;UAS:mCherryNTR larvae across a *foxc1a* and *foxc1b* allelic series (n = 5 per genotype). Yellow dotted lines outline the brain and white dotted lines mark forebrain-midbrain and midbrain-hindbrain boundaries. **(D)** Quantification of dorsal mCherry⁺ meningeal coverage of the midbrain and hindbrain. P-values were calculated using one-way ANOVA with Tukey’s multiple-comparisons test. Error bars represent s.e.m. Unmarked comparisons were not significant (P>0.05). **(E)** Developmental time course (36 hpf–3 dpf) of *foxc1b*:Gal4;UAS:mCherryNTR expression in wild-type and *foxc1a*^-/-^; *foxc1b*^-/-^ embryos (n = 5 per genotype). Yellow dotted lines outline the brain. Asterisks indicate depleted mCherry signal. **(F)** Histological sections of control and mutant larvae stained with Hoechst, showing regions used to quantify mCherry signal. Colored dashed lines indicate quantification zones (n = 5 per genotype). **(G)** Quantification of mCherry signal surrounding the brain from sections in (F). **(H)** RNAscope for *igfbp2a* or *ggctb* with mCherry immunostaining in wild-type and *foxc1a*^-/-^; *foxc1b*^-/-^ larvae (n = 3 per genotype). Arrow indicates an *igfbp2a*⁺ region lacking mCherry signal. P-values were calculated using two-tailed unpaired Student’s t-tests. Error bars represent s.e.m. (’’) = single channels. Scale bars, 100 µm.

To determine the functional requirements of *foxc1a* and *foxc1b*, we generated an allelic series using established mutant alleles (Xu et al., 2018) in combination with the *foxc1b*:Gal4;UAS:mCherryNTR reporter. Loss of *foxc1b* alone did not alter meningeal reporter distribution, as *foxc1a*^+/+^; *foxc1b*^−/−^ and *foxc1a*^+/−^; *foxc1b*^−/−^ larvae appeared phenotypically normal (**Fig. 2C**). In contrast, loss of *foxc1a* resulted in reduced mCherry signal in the caudal hindbrain meninges, and additional loss of *foxc1b* further exacerbated and expanded this phenotype, consistent with a dosage-sensitive requirement for Foxc1 activity (**Fig. 2C**). *foxc1a* mutants develop severe edema and die by 7 dpf, precluding juvenile and adult experiments in compound mutants (Xu et al., 2018).

Quantification of dorsal mCherry signal revealed regional differences in sensitivity to Foxc1 dosage (**Fig. 2D**). Midbrain meninges were comparatively resilient, with significant reductions in mCherry signal detected only in *foxc1a*^−/−^; *foxc1b*^+/−^ and *foxc1a*^−/−^; *foxc1b*^−/−^ mutants. In contrast, caudal hindbrain signal was reduced in *foxc1a*^−/−^; *foxc1b*^+/+^ larvae and further diminished in *foxc1a*^−/−^; *foxc1b*^+/−^ mutants, with little additional loss upon complete *foxc1b* removal. Reduced mCherry signal in *foxc1a*^−/−^; *foxc1b*^−/−^ mutants was evident by 36 hpf and persisted through 72 hpf (**Fig. 2E**). Time-lapse imaging revealed that *fli1a*^+^ mesenchyme emerged readily, whereas mCherry expression initiated dorsally near the rostral hindbrain but failed to expand during the imaging period, with only a few mCherry⁺ cells emerging ventrally toward the caudal hindbrain (**Movie 2**). GFP⁺ and mCherry⁺ cells persisted without widespread morphological evidence of cell loss. Thus, mutant mCherry^+^ meninges fail to encase the brain, suggesting a role for Foxc1 in expanding an initial meningeal progenitor population across the developing brain.

To further resolve regional defects, mCherry⁺ meningeal cells were quantified across five dorsal– ventral domains in sectioned larvae. While dorsal midbrain meninges were largely preserved in *foxc1a*^−/−^; *foxc1b*^−/−^ mutants, all other regions exhibited significant depletion (**Fig. 2F,G**). RNAscope for a broad embryonic meningeal marker *igfbp2a* (**Fig. 1H**) and the leptomeningeal marker *ggctb* was used to assess meningeal identity and differentiation independently of the reporter. In mutants, *igfbp2a* expression was maintained around the brain, including in regions lacking detectable mCherry signal, suggesting *foxc1*-independent activation (**Fig. 2H, Table S1**). In contrast, *ggctb* was restricted to small residual meningeal domains near the midbrain and largely absent from the forebrain and hindbrain, where mCherry loss was most pronounced (**Fig. 2H**). Together, these data demonstrate a conserved, dosage-sensitive requirement for Foxc1 genes during vertebrate meningeal development and reveal regional differences in Foxc1 dependence.

### Pharmacological inhibition identifies pathways required for meningeal formation

To identify developmental pathways required for meningeal establishment, we performed a targeted pharmacological screen using *foxc1b*:Gal4;UAS:mCherryNTR embryos. Inhibitors were applied at 24 hpf, prior to mCherry⁺ meningeal emergence, and embryos were analyzed at 72 hpf, when mCherry⁺ cells encase most of the brain (**Fig. 3A**). Defects were most pronounced in the dorsal hindbrain, the last region to acquire meningeal coverage, which was therefore prioritized for quantification (as illustrated in **Fig. 2C**). Ethanol and DMSO controls did not differ significantly. Inhibition of Hedgehog (Cyclopamine), retinoic acid (DEAB), or FGF (Infigratinib) signaling did not significantly disrupt continuous meningeal coverage, while Notch inhibition (DAPT) caused a modest, non-significant reduction in hindbrain meninges (**Fig. 3B, C**). FGF inhibition did, however, deplete mCherry⁺ meninges at the midbrain-hindbrain interface, likely reflecting its established role in midbrain development (Crossley et al., 1996; Reifers et al., 1998) (**Fig. 3B, Fig. S2A**). In contrast, inhibition of BMP (Dorsomorphin), TGFβ (LY2109761), or Wnt (XAV-939) signaling significantly reduced mCherry⁺ meninges in the dorsal hindbrain (**Fig. 3B, C**). To support a role for Wnt signaling, a second Wnt inhibitor, IWR-1-endo, was applied and similarly reduced hindbrain meninges, although less strongly than XAV-939 (**Fig. 3B, C**). RNAscope at 3 dpf confirmed loss of *foxc1a* transcripts in hindbrain regions depleted of mCherry⁺ meninges, consistent with impaired population of *foxc1*^+^ meningeal cells following pathway inhibition (**Fig. 3D**).

**Figure 3.**
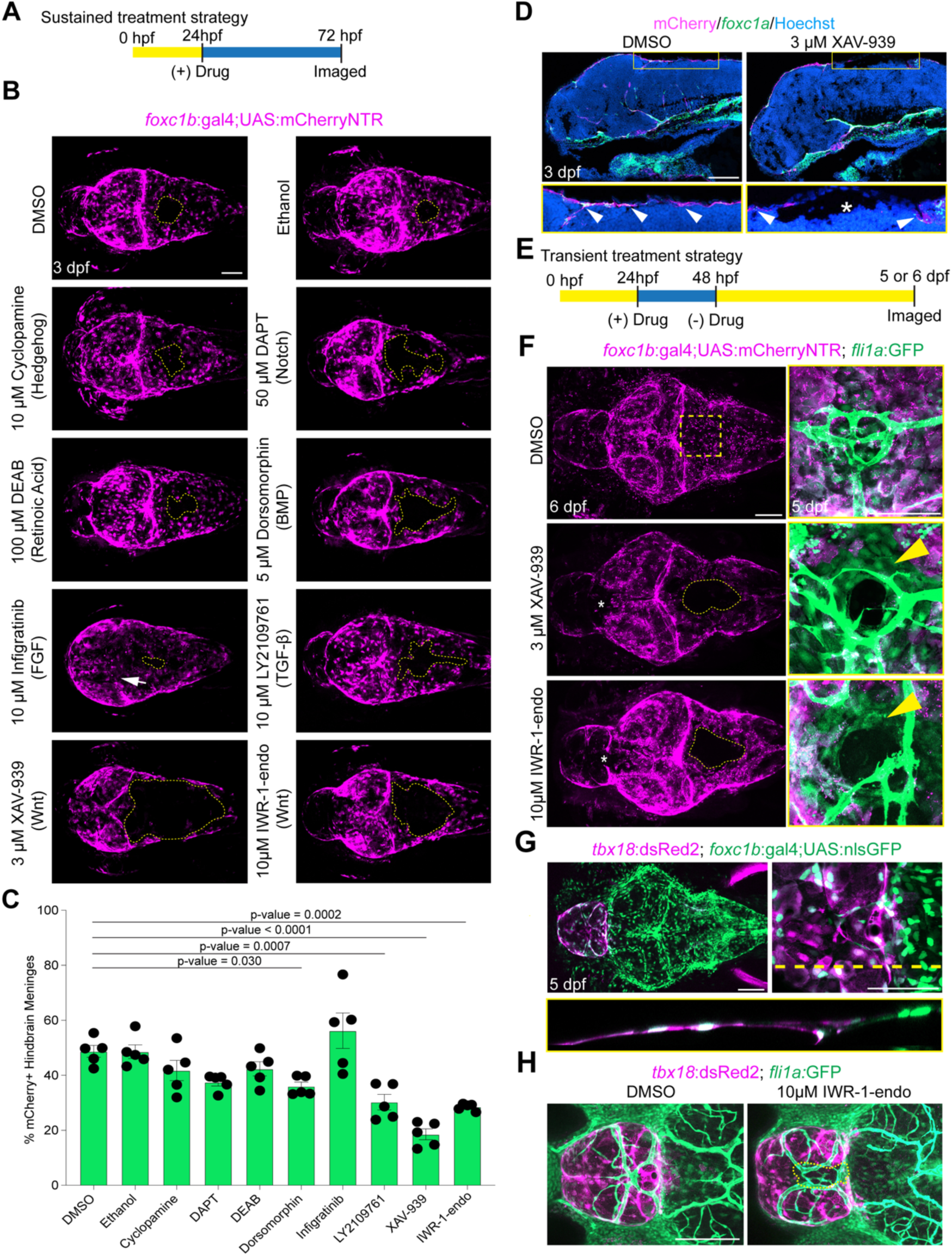
Drug screening of developmental pathways reveals a requirement for Wnt, BMP, and TGFβ signaling during meningeal formation. **(A)** Schematic illustrating the sustained drug treatment strategy. **(B)** Dorsal confocal images of *foxc1b*:Gal4;UAS:mCherryNTR embryos following sustained drug treatment and imaging at 72 hpf (n = 15 embryos per condition). Dotted outlines mark continuous regions of mCherry depletion in the hindbrain. Arrow indicates mCherry depletion at the midbrain-hindbrain interface. **(C)** Quantification of dorsal mCherry⁺ hindbrain meningeal coverage (n = 5 per condition). P-values were calculated using one-way ANOVA with Dunnett’s multiple-comparisons test relative to the DMSO control. Error bars represent s.e.m. Unmarked comparisons were not significant (P>0.05). **(D)** RNAscope for *foxc1a* with mCherry immunostaining in *foxc1b*:Gal4;UAS:mCherryNTR larvae treated with DMSO or XAV-939 (n = 3 per condition). Arrowheads mark *foxc1a*^+^mCherry^+^ regions and asterisks indicate absent signal. Boxes indicate magnified regions. **(E)** Schematic of the transient drug treatment strategy. **(F)** Dorsal confocal images of *foxc1b*:Gal4;UAS:mCherryNTR larvae following transient treatment with DMSO, XAV-939, or IWR-1-endo and imaging at 5 or 6 dpf (n = 10 per condition). Dotted lines mark continuous hindbrain mCherry depletion and asterisks indicate depleted forebrain signal. Box indicates the control region shown at higher magnification with *fli1a*:GFP. Arrowhead indicates *fli1a*:GFP single-positive cells following XAV-939 or IWR-1-endo treatment. **(G)** Dorsal confocal images of 5 dpf *foxc1b*:Gal4;UAS:nlsGFP; *tbx18*:dsRed2 larvae (n = 5). Dorsal (top), magnified (right), and orthogonal views (bottom) show co-expression in the rostral meninges. Dotted lines indicate regions used for orthogonal sections. **(H)** Dorsal-view confocal images of *tbx18*:dsRed2; *fli1a*:GFP larvae following transient treatment with DMSO or IWR-1-endo and imaging at 6 dpf. Dotted outline marks disruption of dsRed2⁺ cells at the forebrain-midbrain interface (n = 5 larvae per condition). Scale bars, 100 µm.

To determine whether transient Wnt or BMP inhibition was sufficient to disrupt meningeal development, embryos were treated with XAV-939, IWR-1-endo, or Dorsomorphin from 24–48 hpf and analyzed at 5–6 dpf (**Fig. 3E**). Treated larvae were comparable in size to controls but exhibited persistent depletion of mCherry signal in the medial hindbrain meninges for all treatments (**Fig. 3F, Fig. S2C**). In *foxc1b*:Gal4;UAS:mCherryNTR; *fli1a*:GFP larvae, GFP^+^ hindbrain mesenchymal cells persisted following XAV-939 and IWR-1-endo treatment, but mCherry reporter activation was selectively impaired (**Fig. 3F, Table S1**). In contrast, Dorsomorphin treatment depleted both mCherry and GFP signal, consistent with broader loss of mesenchymal cells (**Fig. S2C**). RNAscope at 6 dpf confirmed selective loss of *foxc1a*⁺*/*mCherry⁺ hindbrain meninges following IWR-1-endo treatment (**Fig. S2C**).

The *foxc1b*:Gal4;UAS:mCherryNTR reporter generally displayed reduced baseline signal in the medial-dorsal forebrain (**Fig. 1A-D**), but XAV-939 and IWR-1-endo treatment suggested further depletion of mCherry⁺ meninges (**Fig. 3F**). To further assess this phenotype, we examined larvae carrying the forebrain meningeal reporter *tbx18*:dsRed2, which co-expresses with *foxc1b*:Gal4;UAS:nlsGFP (**Fig. 3G, Table S1**). Transient IWR-1-endo treatment (**Fig. 3E**) of *tbx18*:dsRed2; *fli1a*:GFP embryos disrupted dsRed2 signal, particularly at the forebrain–midbrain boundary, indicating that Wnt signaling is required for proper establishment of forebrain meninges (**Fig. 3H**). Because *Tbx18* also labels mural cells (DeSisto et al., 2020; Guimaraes-Camboa et al., 2017), co-imaging of *tbx18*:dsRed2 and *fli1a*:GFP confirmed that dsRed2 depletion was not solely attributed to loss of vessels and associated mural cells, although vessel architecture was abnormal (**Fig. 3H**). Together, these data identify TGFβ, BMP and Wnt signaling as regulators of meningeal development, with Wnt signaling required for proper establishment of both hindbrain and forebrain meninges.

### Adult zebrafish meninges are comprised of transcriptionally distinct fibroblasts

During embryonic stages, zebrafish meninges appear as a single cellular layer (**Fig. 1K**), whereas adult zebrafish meninges are multilayered (Galanternik et al., 2025). To define adult meningeal populations, we performed single-nucleus multiomic profiling.

In adult *foxc1b*:Gal4;UAS:mCherryNTR fish (≥3 months and ≥20 mm), robust mCherry signal surrounded the brain and lined the inner surface of the skull, with enrichment in the midbrain and hindbrain (**Fig. 4A**). Histology confirmed reduced dorsal forebrain mCherry signal, while *tbx18* transcripts and *tbx18*:dsRed2 expression persisted in this region, consistent with regional differences in meningeal composition or the presence of specialized structures such as the tela choroidea (**Fig. S3A-C**). Histological analysis of *sox10*;ubi:loxP-EGFP-loxP-mCherry adults (from embryos treated with tamoxifen at 1 dpf) demonstrated neural crest contributions to the forebrain meninges, consistent with lineage boundaries described for the overlying calvarial bones (**Fig. S3D**) (Kague et al., 2012; Mongera et al., 2013).

**Figure 4.**
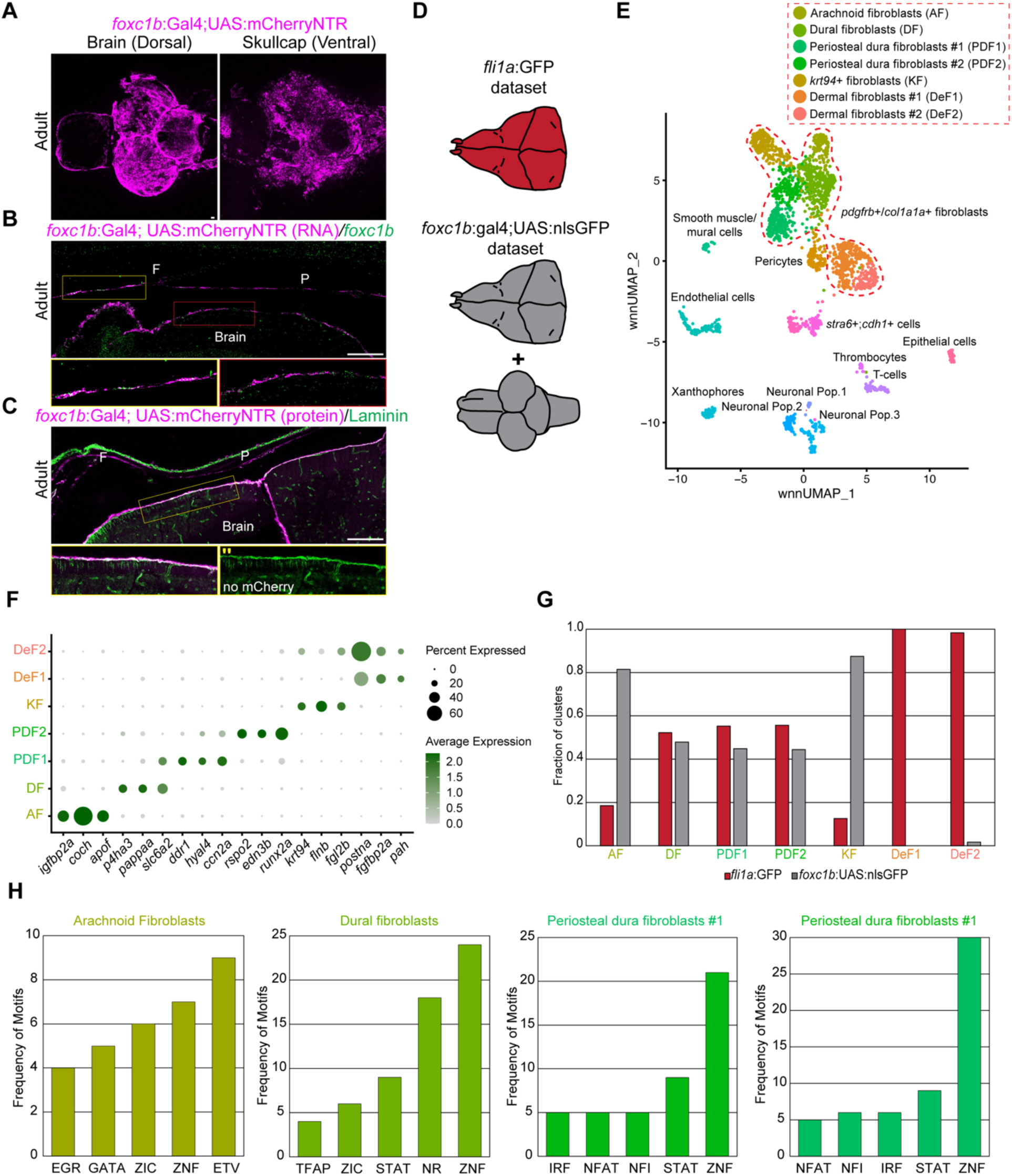
Single-cell genomics reveals multiple fibroblast cell types within zebrafish meninges. **(A)** Dorsal view of a dissected brain and ventral view of a dissected skullcap from adult *foxc1b*:Gal4;UAS:mCherryNTR fish, showing meningeal mCherry expression (n = 5). **(B)** RNAscope for *foxc1b* and mCherry in adult *foxc1b*:Gal4;UAS:mCherryNTR fish (n = 3). **(C)** Laminin and mCherry immunostaining in adult *foxc1b*:Gal4;UAS:mCherryNTR fish (n = 3). **(D)** Schematic of transgenic lines and tissues used for single-cell genomic analyses. **(E)** UMAP of merged datasets with cluster annotations. Red dashed lines delineate fibroblast clusters, with predicted identities indicated within the red box. **(F)** Dot plot of marker genes enriched in fibroblast clusters. **(G)** Fraction of each fibroblast cluster derived from each dataset, with colors corresponding to (D). **(H)** Frequency of enriched transcription factor family motifs within representative clusters. F= frontal bone, P= parietal bone. Boxes indicate magnified regions. (’’) = single channels. Scale bars, 100 µm.

RNA analysis confirmed co-detection of mCherry and *foxc1b* transcripts in meninges associated with both brain and skull (**Fig. 4B**). Pan-laminin immunostaining selectively marked mCherry⁺ cells associated with the brain but not those adjacent to the skull, suggesting molecular heterogeneity within adult meninges (**Fig. 4C**).

To define transcriptional and epigenetic profiles of adult meningeal populations, connective tissues associated with the brain or skull were isolated from *foxc1b*:Gal4;UAS:nlsGFP and *fli1a*:GFP fish, which we previously reported in dura mater (Farmer et al., 2024), and subjected to multiome analysis (**Fig. 4D**). After quality control, 1,224 cells from the *fli1a*:GFP dataset and 1,188 cells from the *foxc1b*:Gal4;UAS:nlsGFP dataset were recovered. Unsupervised clustering of merged datasets identified 17 populations (**Fig. 4E, Fig. S4A, Table S2**), including mesenchymal populations expressing connective tissue markers *pdgfrb* and *col1a1a* (**Fig. S4B**), as well as pericytes, smooth muscle/mural cells, endothelial cells, epithelial cells, immune populations, pigment cells, and neuronal cell types (**Fig. 4E, Fig. S4C**). Relative abundance of non-mesenchymal populations differed modestly between datasets, whereas endothelial, epithelial, and T cell populations were present at comparable levels (**Fig. S4D)**.

Within mesenchymal populations, dermal fibroblasts were enriched in the *fli1a*:GFP dataset, while *krt94*⁺ mesenchymal cells, captured in our prior datasets (Farmer et al., 2024) and validated to be a connective tissue bordering the skull (**Fig. S4E**), were enriched in the *foxc1b*:Gal4;UAS:nlsGFP dataset (**Fig. 4F, G**). With the exception of arachnoid fibroblasts, remaining mesenchymal populations showed comparable contributions from both datasets (**Fig. 4F, G**). Analysis of cluster-enriched genes identified previously uncharacterized fibroblast populations, including arachnoid fibroblasts, dural fibroblasts, and two transcriptionally distinct periosteal dura fibroblast populations that were further validated in this study (**Fig. 4E-G**).

To relate zebrafish meningeal populations to mammalian counterparts, we reanalyzed published adult mouse leptomeningeal and dura mater single-cell datasets (Pietila et al., 2023) and performed cross-species comparisons using module scores derived from zebrafish cluster-enriched genes and their mouse paralogs (**Fig. S5A–D**). The *coch*⁺ cluster corresponded most closely to mouse pial fibroblasts and inner arachnoid cells (BFB1/BFB2), whereas the *slc6a2*⁺ cluster matched a previously undescribed dura population enriched for *Pappa* (**Fig. S5E-F**). The *ccn2a*⁺ periosteal dura fibroblast cluster aligned with mouse outer dura populations expressing *Matn4* and *Nppc* (Farmer et al., 2021) (**Fig. S5G**). In contrast, the *rspo2*⁺ periosteal dura fibroblast cluster lacked clear correspondence to known mouse meningeal populations (**Fig. S5H**).

We next investigated the presence of arachnoid barrier cells (ABCs) using markers identified in adult mice (Pietila et al., 2023), which were not strongly co-enriched within any zebrafish cluster (**Fig. S6A**). Although an undescribed *cdh1*⁺ population was previously annotated as *stra6*⁺;*cdh1*⁺, histological analysis of *cdh1*:mlanYFP did not identify YFP⁺ cells associated with the brain or inner surface of the skull, and RNAscope did not identify a specifically enriched *stra6*⁺ population outside the skin, despite low expression in the suture and brain (**Fig. S6B-C**). Weak *dpp4*, *crabp2a*, and *cldn11a* expression was detected within putative dural/periosteal populations, but a definitive ABC homolog remains unclear (**Fig. S6A**).

Analysis of chromatin accessibility revealed cluster-specific regulatory programs. Arachnoid fibroblasts were enriched for ETV, EGR, and GATA motifs, dural fibroblasts for TFAP, STAT, and nuclear receptor motifs, and both populations for ZIC family motifs, consistent with known roles in meningeal biology (Inoue et al., 2008) (**Fig. 4H**). Periosteal dura fibroblast clusters shared enrichment for IRF, NFAT, and NFI motifs, while all meningeal populations exhibited enrichment for ZNF family motifs (**Fig. 4H**).

To spatially localize transcriptionally defined populations, RNAscope was performed in adult zebrafish. *ggctb*⁺*igfbp2a*⁺ cells, also observed at larval stages, persisted adjacent to the brain but were largely absent beneath the skull, despite poor representation in single-cell datasets and co-expression of *ggctb* and *foxc1b*:Gal4;UAS:GFP (**Fig. 5A; Fig. S7A, Table S1**). Cells adjacent to the brain also expressed transcripts encoding the secreted chemokine *cxcl12a* and the retinoic acid–synthesizing enzyme *aldh1a2* (**Fig. S7B**). Arachnoid cells co-expressing *coch* and *igfbp2a* formed a discrete population superficial to *ggctb*⁺ cells (**Fig. 5B,C, Table S1**). In contrast, dural populations marked by *slc6a2* and periosteal dura fibroblasts marked by *ccn2a* and *rspo2* localized preferentially to the skull-associated side of the meninges **(Fig. 5D,E, Table S1**). *ccn2a*⁺ and *rspo2*⁺ cells were intermingled within a single bone-adjacent layer, consistent with transcriptionally distinct periosteal dura fibroblast populations occupying the same anatomical compartment.

**Figure 5.**
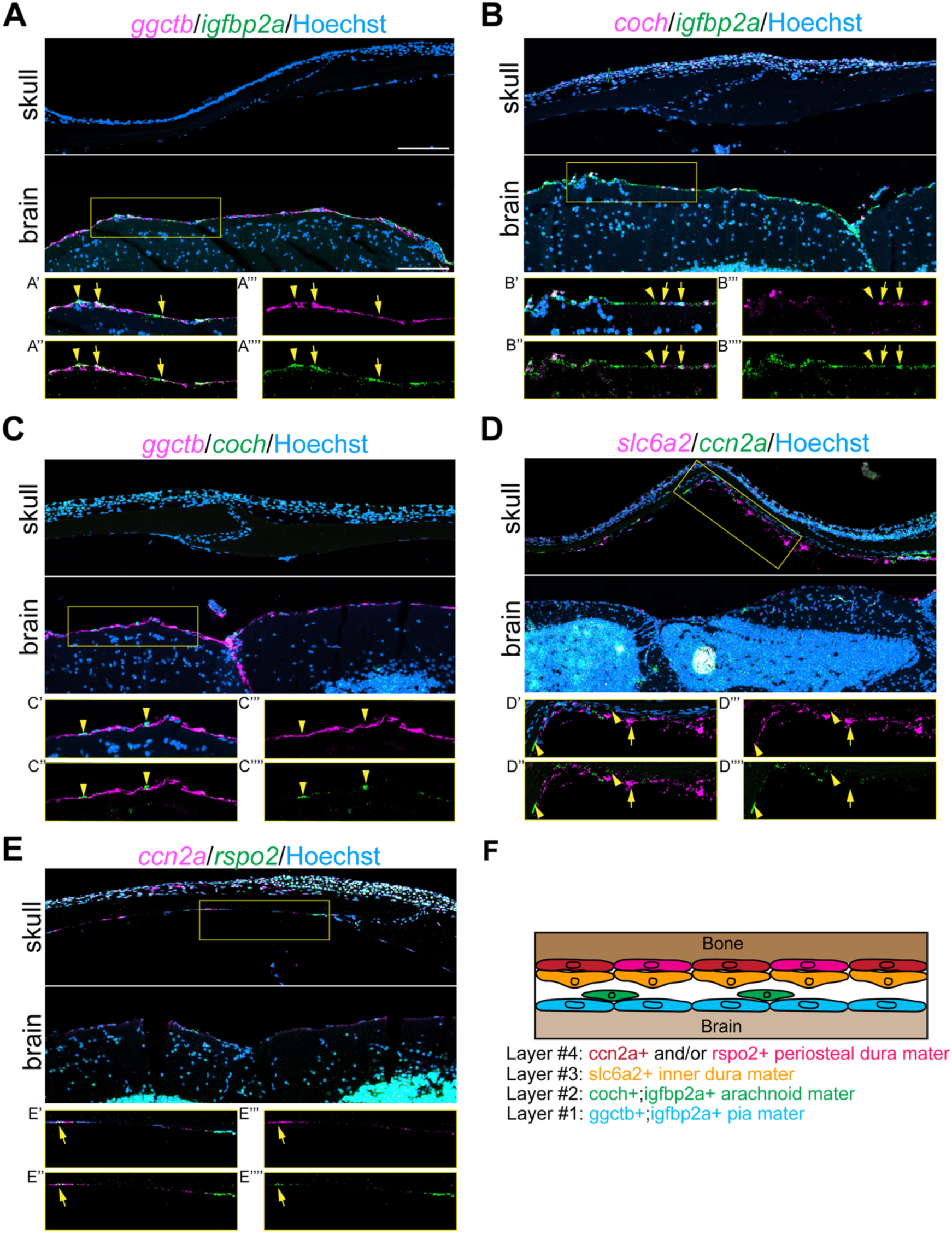
Meningeal fibroblasts are spatially organized into distinct layers between the skull and brain in the adult zebrafish. **(A)** RNAscope for *ggctb* and *igfbp2a* shows double-positive cells surrounding the brain (arrows), with interspersed *igfbp2a*⁺ cells (arrowhead). **(B)** RNAscope for *coch* and *igfbp2a* shows scattered double-positive cells around the brain (arrows) and *igfbp2a*⁺ cells along the brain surface (arrowhead). **(C)** RNAscope for *ggctb* and *coch* reveals non-overlapping domains, with *coch*⁺ cells (arrowheads) superficial to *ggctb*⁺ cells. **(D)** RNAscope for *slc6a2* and *ccn2a* shows *ccn2a*^high^ cells (arrowheads) closest to the bone and *slc6a2*^high^ cells (arrows) farther from the bone. Low *slc6a2* expression is also detected around the brain. **(E)** RNAscope for *ccn2a* and *rspo2* shows both genes within the same bone-adjacent layer, including double-positive cells (arrow). Both are absent around the brain. **(F)** Model of meningeal subtype organization in the adult zebrafish skull. Boxes indicate magnified regions. (’) = magnified region, (’’) = no Hoechst, (’’’) and (’’’’) = single channels. All stains were performed in biological triplicates. Scale bars, 100 µm.

Together, these data demonstrate that adult zebrafish meninges comprise a multilayered and transcriptionally heterogeneous structure consisting of at least four distinct populations that we tentatively define by their spatial organization and/or transcriptional profiles: a pial layer (*ggctb*⁺*igfbp2a*⁺), a discrete arachnoid layer (*coch*⁺), an inner dura mater layer (*slc6a2*⁺), and a periosteal dura layer composed of intermingled *ccn2a*⁺ and *rspo2*⁺ fibroblasts (**Fig. 5F**).

### Larval meningeal ablation reveals long-term consequences on calvarial development

To assess the functional requirements for meninges during development, we ablated mCherry⁺ cells using *foxc1b*:Gal4;UAS:mCherryNTR. Because juvenile or adult treatments resulted in lethality within 24 hours, subsequent analyses focused on larval ablation. Overnight metronidazole (MTZ) treatment initiated at 5 dpf resulted in widespread loss of mCherry signal, confirming efficient ablation (**Fig. 6**, **Fig. S8A,B**). Consistent with loss of meningeal integrity, expression of the broad meningeal marker *igfbp2a* was markedly reduced around the brain at 0 and 1 days post-treatment (dpt), with discontinuous recovery by 2 dpt, indicating partial regeneration of the meninges (**Fig. S8C**). Analysis of *foxc1b*:Gal4;UAS:mCherryNTR; UAS:GFP larvae revealed recovery of GFP⁺/*igfbp2a*⁺ cells by 2 dpt, consistent with meningeal re-establishment despite persistent loss of mCherry reporter expression (**Fig. S8D**). MTZ-treated larvae exhibited reduced long-term survival, with 54% surviving to 18 dpt compared to 87% of DMSO-treated controls (**Fig. S8E**).

**Figure 6.**
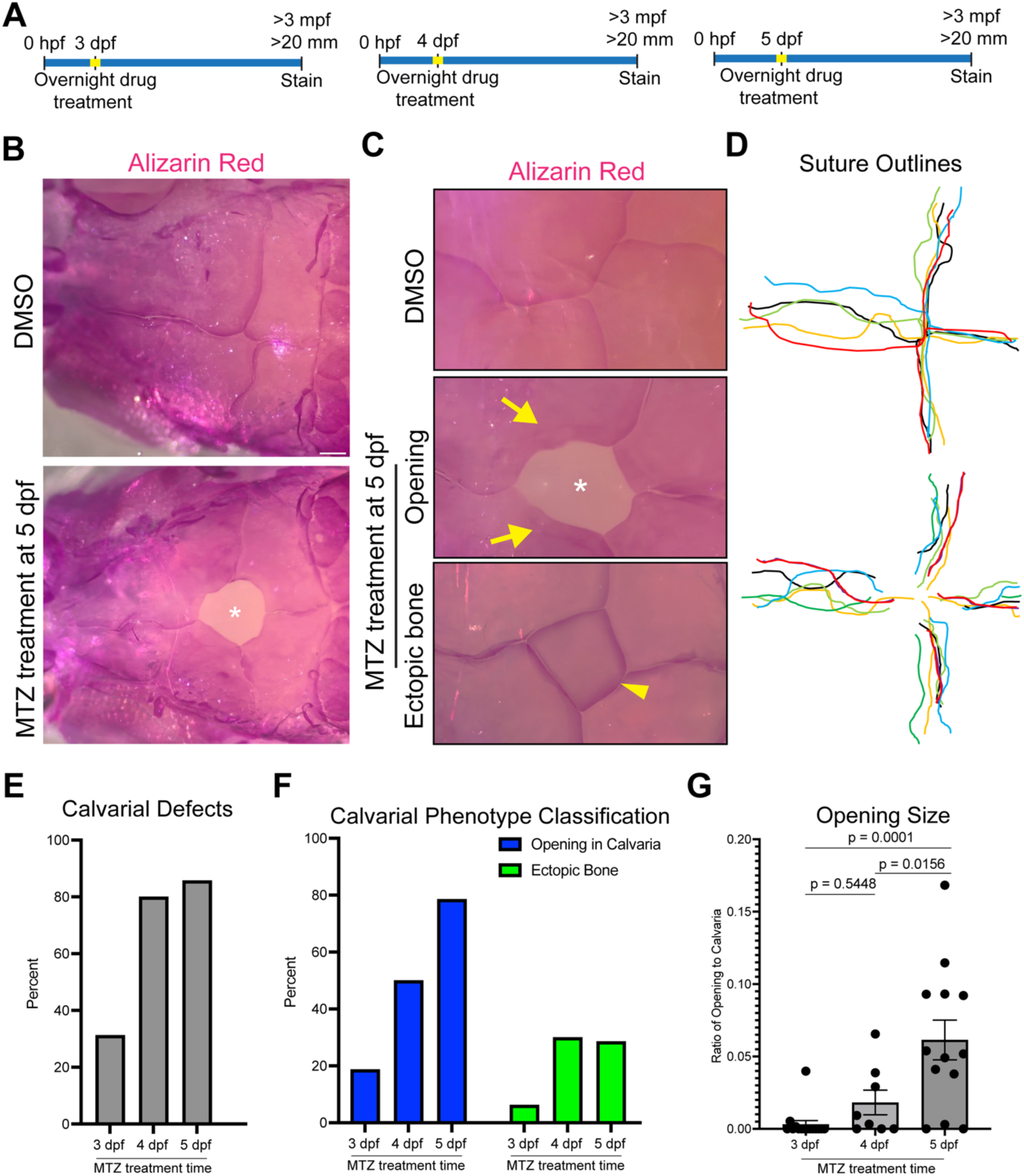
Larval ablation of meninges leads to persistent calvarial defects. **(A)** Schematic of MTZ treatment strategies. **(B)** Dorsal views of Alizarin-stained skulls from DMSO- and MTZ-treated adult zebrafish (n = 13 fish per condition). **(C)** Representative bregma regions showing the range of calvarial phenotypes following MTZ treatment. Arrows indicate irregular bone borders suggestive of misdirected growth, arrowhead indicates ectopic bone, and asterisks mark calvarial openings. **(D)** Hand-drawn outlines of suture borders in control and MTZ-treated zebrafish. **(E)** Quantification of MTZ-treated zebrafish exhibiting calvarial defects following ablation at different developmental stages. **(F)** Quantification of calvarial phenotype classes by treatment stage. **(G)** Quantification of calvarial openings as the ratio of opening area to total calvarial area. P-values were calculated using one-way ANOVA with Tukey’s multiple-comparisons test. Error bars represent s.e.m. Scale bars, 100 µm.

To test for long-term consequences of treatment, MTZ-treated fish were raised to adulthood (≥3 months old and ≥20 mm) and their calvaria were analyzed. Alizarin Red staining revealed widespread calvarial defects, with the most prevalent being a persistent opening in the skull (**Fig. 6B, C**). These openings were frequently associated with misdirected bone growth, including bone folding and growth away from the opening, while ectopic bone occasionally formed between the frontal and parietal bones (**Fig. 6C**). Defects were reproducibly localized to the bregma, where cranial sutures converge (**Fig. 6D**).

To determine whether calvarial phenotypes were sensitive to the timing of meningeal ablation, MTZ treatment was initiated at 3, 4, or 5 dpf. Ablation at 3 dpf, when the brain is nearly encased by meninges, resulted in the lowest incidence of calvarial defects (∼33%), compared with ∼80% following ablation at 4 or 5 dpf (**Fig. 6E**). Later ablation increased the frequency of calvarial openings and ectopic bone formation (**Fig. 6F**). Ablation at 5 dpf also increased opening size compared with ablation at 3 dpf and 4 dpf (**Fig. 6G**). These findings demonstrate stage-dependent sensitivity of calvarial development to meningeal disruption.

To investigate the basis of calvarial openings, osteogenesis was examined during skull development beginning at 8 mm standard length, which healthy zebrafish typically reach at approximately 3–4 weeks under our laboratory conditions. At this stage, before calvarial bones encase the brain (Topczewska et al., 2016), pigmented dural cells (Galanternik et al., 2025) normally populate the head, and uniformly cover the midbrain and hindbrain in controls (**Fig. 7A**). In contrast, MTZ-treated fish exhibited pronounced depletion of pigmented dural cells, consistent with meningeal disruption preceding cranial suture formation and a failure to re-establish an appropriate environment for normal pigment cell repopulation (**Fig. 7A**).

**Figure 7.**
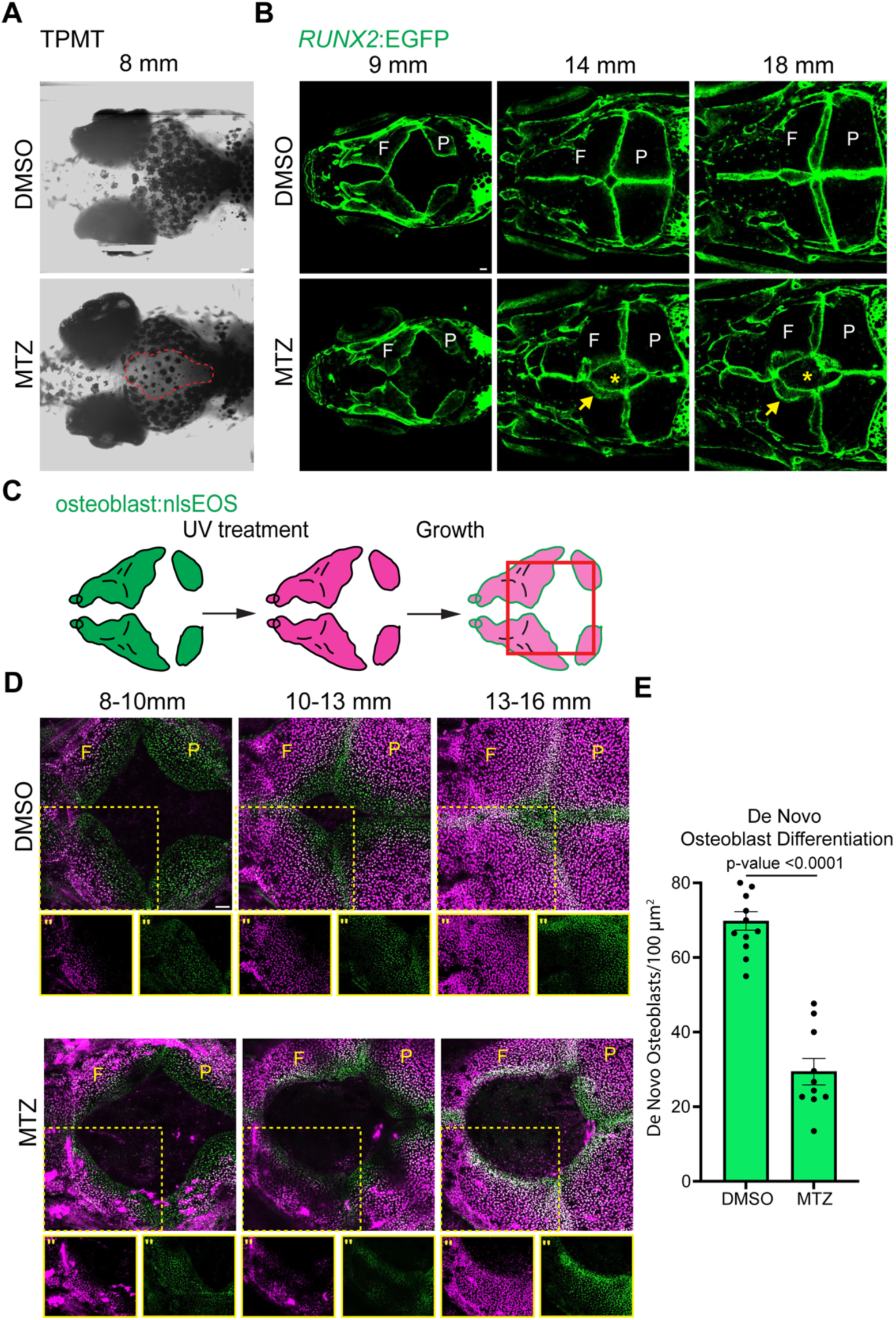
Osteogenesis is disrupted by early ablation of meninges. **(A)** Transmitted light images of control- and MTZ-treated zebrafish at 8 mm (n = 5 fish per condition). Dotted line demarcates the region of pigment depletion. **(B)** Dorsal confocal images from serial imaging of *foxc1b*:Gal4;UAS:mCherryNTR (not imaged); *RUNX2*:GFP zebrafish treated with DMSO or MTZ from 9–18 mm (n = 6 per condition). Arrows indicate abnormal overlapping bone and asterisks mark RUNX2-negative calvarial openings. **(C)** Schematic of the treatment and imaging strategy used to quantify *de novo* osteoblast formation. Red box indicates the region imaged in subsequent panels. **(D)** Serial confocal imaging of DMSO- and MTZ-treated osteoblast:nlsEOS fish. Fish were sequentially photoconverted at approximately 8, 10, and 13 mm and imaged at 10, 13, and 16 mm, respectively (n = 5 per condition). Dotted boxes indicate regions shown at higher magnification. F = frontal bone, P = parietal bone. **(E)** Quantification of de novo osteoblasts at bone fronts. P-values were calculated using two-tailed unpaired Student’s t-tests. Error bars represent s.e.m. (’’) = single channels. Scale bars, 100 µm.

Repeated live imaging of *RUNX2*:EGFP fish revealed no differences between control and MTZ-treated animals at 9 mm, when calvarial bones have initiated growth but remain separated (**Fig. 7B**). By 14 mm, when bones normally approximate, MTZ-treated fish exhibited persistent openings at the prospective bregma, with *RUNX2*⁺ bone fronts failing to converge and instead folding or growing over adjacent bone. These defects persisted through 18 mm, when cranial sutures are normally established. To assess osteoblast differentiation, we used the osteoblast:nlsEOS reporter to track *de novo* osteoblast formation (Farmer et al., 2024) (**Fig. 7C**). In controls, newly differentiated osteoblasts accumulated as bone fronts approached and decreased after bones met (**Fig. 7D**). In contrast, MTZ-treated fish showed a marked reduction in de novo osteoblast differentiation at open bone fronts, with fewer newly-differentiated osteoblasts detected near calvarial openings (**Fig. 7D, E**).

Given the reproducible localization of defects at the bregma, we tested whether focal meningeal ablation was sufficient to induce long-term craniofacial abnormalities. Targeted two-photon ablation of meningeal cells at the dorsal midbrain–hindbrain boundary resulted in efficient but transient loss of meningeal fluorescence, with robust regeneration by 30 hours post-ablation (**Fig. S9A**). Fish subjected to focal ablation and raised to adulthood exhibited no craniofacial abnormalities or calvarial openings (**Fig. S9 B**). These findings suggest that large-scale, rather than focal, meningeal disruption is required to compromise calvarial development.

### Impaired meningeal regeneration is associated with calvarial and brain defects

To define tissue-level disruptions associated with impaired osteogenesis and calvarial openings, we performed histological and whole-mount analyses of DMSO- and MTZ-treated skulls. Plastic embedding preserved skull-brain architecture and revealed the brain closely apposed to the skull with intact intervening tissues in controls (**Fig. 8A**). In contrast, MTZ-treated skulls failed to fully encase the brain, which adhered to the overlying dermis and exhibited disorganization of underlying neural tissue (**Fig. 8A**).

**Figure 8.**
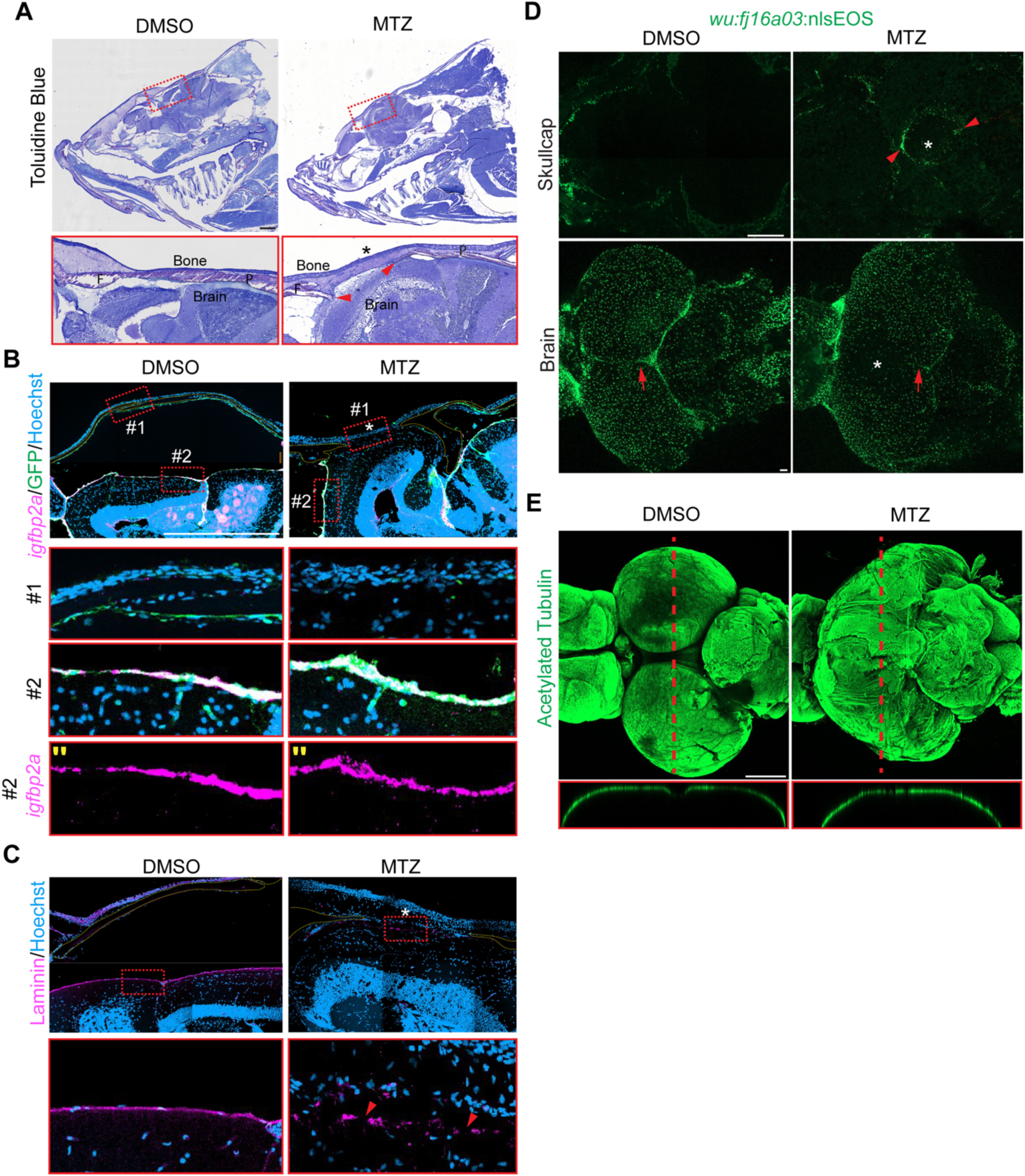
Impaired dorsal meningeal regeneration is associated with irregular calvarial and brain morphology. **(A)** Toluidine blue staining of plastic-embedded heads from DMSO- and MTZ-treated fish (n = 3 fish per condition). Arrowheads mark bone ends in MTZ-treated skull. **(B)** RNAscope for *igfbp2a* with GFP immunostaining in *foxc1b*:Gal4; UAS:mCherryNTR;UAS:GFP heads (n = 4 fish per condition). **(C)** Laminin immunostaining of DMSO- and MTZ-treated heads (n = 3 fish per condition). **(D)** Confocal images of dissected skullcaps and brains from DMSO- and MTZ-treated *foxc1b:*Gal4;UAS:mCherryNTR (not imaged); *wu:fj16a03*:nlsEOS fish (n = 3 fish per condition). Arrowheads mark EOS⁺ cells adherent to the skullcap. Arrows indicate EOS⁺ signal between the optic tecta. **(E)** Confocal imaging of whole brains stained for acetylated tubulin (n = 3 fish per condition). Red dotted lines indicate regions used for orthogonal sections. Yellow dotted outlines mark bones and asterisks mark calvarial openings. Dotted boxes indicate regions shown at higher magnification. F= frontal bone, P= parietal bone. (’’) = single channels. Scale bars, 100 µm.

To assess meningeal regeneration at calvarial openings, we performed RNAscope in *foxc1b*:Gal4;UAS:mCherryNTR;UAS:GFP animals, as mCherry expression does not recover after MTZ treatment (**Fig. S8D**). In controls, GFP⁺ meningeal cells lined the ventral skull surface and *igfbp2a*⁺GFP⁺ cells encased the brain (**Fig. 8B**). In MTZ-treated tissue, brain tissue remained adherent to the skin at calvarial openings where neither GFP nor *igfbp2a* signal was detected, despite persistence of GFP⁺ cells adjacent to surrounding bone and *igfbp2a*⁺ GFP⁺ meninges away from the opening (**Fig. 8B**). These data indicate impaired meningeal regeneration specifically at sites of failed bone formation. Consistent with previous associations between meningeal disruption and basement membrane defects (Inoue et al., 2008; Zarbalis et al., 2007), laminin was continuous around the brain in controls but disorganized in MTZ-treated animals (**Fig. 8C**).

Because some MTZ-treated skulls exhibited ectopic bone, we examined meningeal regeneration at these sites. RNAscope revealed irregular brain–skull adhesion and absent *igfbp2a* expression beneath ectopic bone, indicating that bone can form without normal meningeal repopulation but is associated with aberrant calvarial patterning (**Fig. S10**).

Consistent with histological and molecular analyses, imaging of the *wu:fj16a03*:nlsEOS reporter that labels brain-associated meningeal cells (**Fig. S11**) revealed depletion of EOS⁺ cells on the surface of brain overlying calvarial openings and enrichment along the periphery of the defect on the skullcap (**Fig. 8D**). Single-cell transcriptomic analysis of DMSO- and MTZ-treated skullcaps revealed largely comparable cellular compositions, except for a cluster enriched in MTZ-treated samples expressing markers characteristic of arachnoid fibroblasts normally associated with brain meninges (**Fig. S12A-D; Table S3**).

In controls, bilateral optic tecta were clearly demarcated by *wu:fj16a03*:nlsEOS⁺ cells, whereas this organization was lost in MTZ-treated brains (**Fig. 8D**). Whole-mount staining for acetylated tubulin confirmed disrupted brain architecture, including loss of separation between the optic tecta (**Fig. 8E**). Together, these findings demonstrate that impaired meningeal regeneration is associated with persistent defects in skull and brain development, supporting a role for the meninges in coordinating the development of adjacent tissues.

## Discussion

The meninges are critical connective tissues supporting the brain and skull (Dasgupta and Jeong, 2019), yet their cellular composition, development, and signaling functions remain incompletely understood. Here, we define the developmental emergence, molecular heterogeneity, and functional importance of zebrafish meningeal populations across larval and adult stages. Integrating genetic, genomic, and functional approaches, we identify conserved developmental programs and roles for the meninges in calvarial growth and brain organization.

### Developmental emergence and heterogeneity of zebrafish meninges

Using an enriched *foxc1b*:Gal4 reporter, we show that zebrafish meninges emerge in a spatiotemporally ordered manner, with rostral meninges preceding caudal populations. Although the progression of *Foxc1* expression across mammalian meninges has not been described in similar detail, other differentiation patterns show similarities. In zebrafish, the leptomeningeal marker *ggctb* emerges prominently ventrally before expanding dorsally. Similarly, Raldh2^+^ cells in the murine meninges first appear ventrally and progress in a lateral-to-medial manner to cover the brain (Siegenthaler et al., 2009). Whether these patterns reflect conserved sequences of meningeal differentiation and stratification will require future analysis with comparable markers (DeSisto et al., 2020; Siegenthaler et al., 2009; Vivatbutsiri et al., 2008). At larval stages, fibroblast and leptomeningeal markers are co-expressed within a single cellular layer, while progressive leptomeninges marker expression suggests increasing differentiation. When stratification emerges remains unclear, but the transition to fully stratified meninges may occur later in zebrafish than in mice, paralleling the later emergence of zebrafish calvarial bones (Topczewska et al., 2016).

In adults, single-cell profiling and spatial validation revealed a multilayered meningeal architecture comprising at least four transcriptionally and spatially distinct populations. These findings provide molecular resolution to prior anatomical descriptions (Galanternik et al., 2025) and demonstrate that, as in mammals, zebrafish meninges comprise specialized layers with distinct identities. Specialized populations such as arachnoid barrier and dural border cells lacked clear zebrafish counterparts, suggesting potential differences in meningeal organization between species that warrant future investigation. Thus, zebrafish provide a tractable model for dissecting conserved mechanisms of meningeal stratification while also revealing potential evolutionary differences in meningeal complexity. Future studies will be needed to define the conserved functional requirements of each meningeal cell type.

### Conserved regulatory programs governing meningeal development

Functional analyses demonstrate a conserved, dosage-sensitive requirement for Foxc1 during meningeal development, with regional differences in sensitivity. Loss of *foxc1a* preferentially affected hindbrain meninges, while additional loss of *foxc1b* exacerbated defects, consistent with mammalian studies linking FOXC1 dosage to meningeal integrity (Aldinger et al., 2009; Siegenthaler et al., 2009; Zarbalis et al., 2007). Persistence of some meningeal populations in *foxc1*-deficient contexts suggests that additional transcriptional regulators contribute to meningeal specification. Although both paralogs show comparable meningeal expression, *foxc1a* appears more critical, consistent with asymmetric requirements in other zebrafish tissues, where *foxc1a* loss produces stronger phenotypes while *foxc1b* contributes when *foxc1a* activity is reduced (Ferre-Fernandez et al., 2020; Topczewska et al., 2001). Thus, the paralogs provide overlapping but unequal contributions to Foxc1 activity, although whether this reflects differences in regulation or intrinsic protein activity remains unclear.

Pharmacological perturbation identified TGFβ, BMP and Wnt signaling as regulators of meningeal establishment, particularly for late-forming hindbrain populations. Wnt inhibition impaired meninges establishment without broadly affecting mesenchymal survival, consistent with previous evidence that Wnt regulates meningeal expansion in mice (Choe et al., 2014). In contrast, BMP inhibition depleted both mesenchymal and meningeal-enriched markers, suggesting distinct requirements for these pathways. Wnt inhibition also depleted *tbx18*:dsRed2⁺ forebrain meninges, although effects on other *tbx18*⁺ populations, including mural cells, cannot be excluded. TGFβ inhibition similarly disrupted early meningeal development but requires further investigation. Motif enrichment analyses further implicate transcription factor families previously associated with meningeal biology, including ZIC factors, suggesting that layered combinations of signaling inputs and transcriptional programs generate meningeal diversity. The identification of population-specific enrichment of GATA, TFAP, and NFAT motifs highlights new potential regulatory pathways for future study.

### Meninges as regulators of calvarial osteogenesis

A central finding of this study is that meningeal integrity supports proper calvarial development. Larval meningeal ablation resulted in persistent calvarial openings, misdirected and ectopic bone growth, and reduced *de novo* osteoblast differentiation at bone fronts, despite apparently normal initial bone formation. Focal ablation was insufficient to produce long-term defects, suggesting that large-scale or sustained meningeal disruption is required to compromise calvarial patterning. Together with ectopic bone formation in regions lacking canonical meningeal markers, these findings suggest that meninges provide positional or organizational cues supporting bone growth rather than acting solely as inductive sources of osteogenic signals. Roles for meninges in maintaining suture patency have been described in mammals (Cooper et al., 2012; Greenwald et al., 2000; Kwan et al., 2008; Opperman et al., 1993; Slater et al., 2008), and hypomorphic *Foxc1* mutations produce comparable calvarial openings (Zarbalis et al., 2007). Aberrant adhesion of the meninges to adjacent tissues may help re-establish a closed CSF compartment, but at the expense of an appropriate local environment for brain and bone development.

### Meningeal regeneration and tissue coordination

Despite partial regeneration following ablation, meninges fail to re-establish normal tissue architecture at calvarial openings. The increased severity of calvarial phenotypes following ablation at 5 dpf compared with 3 dpf coincides with expanded expression of the leptomeningeal marker ggctb and may reflect progressive meningeal differentiation that restricts regenerative capacity. Loss of meningeal markers, abnormal brain–dermis adhesion, and tissue disorganization are associated with altered brain morphology, including impaired separation of the optic tecta. Together, these findings highlight the meninges as an important interface coordinating brain and skull development.

Consistent with mouse studies, meningeal loss may disrupt delivery of key secreted factors, such as retinoic acid (Como et al., 2025; Haushalter et al., 2017; Matrongolo et al., 2023; Mishra et al., 2016; Siegenthaler et al., 2009) and CXCL12 (Borrell and Marin, 2006), both of which regulate neural development and were also detected in zebrafish meninges adjacent to brain tissue. The pronounced brain phenotypes presented here support conserved meningeal–neural interactions across vertebrates.

### Limitations

1. Technical limitations may have impeded capture of pial fibroblasts closely associated with the brain and potentially other meningeal populations, limiting our ability to fully resolve meningeal heterogeneity and assess relationships between larval and adult *ggctb*⁺ cells. Future spatial genomics approaches will help resolve these populations and their developmental relationships.
2. *foxc1a/b* function in multiple developmental processes, including mural cell differentiation, somitogenesis, cardiac and craniofacial development, and their combined loss is associated with hydrocephalus (Chrystal et al., 2021; Ferre-Fernandez et al., 2020; Li et al., 2015; Whitesell et al., 2019; Xu et al., 2018). Although our findings support conserved roles for Foxc1 in meningeal development, conditional approaches will be needed to define tissue-specific requirements within the meninges (Shin et al., 2023).
3. Whole-embryo pharmacological perturbations limit the cellular and temporal resolution of the signaling requirements identified here. Gal4/UAS and inducible Cre-based approaches will enable more precise analysis of these pathways.
4. Although *foxc1b*:Gal4;UAS:mCherryNTR labels a restricted population, expression is not exclusive to the meninges, and ablation of other labeled populations may contribute to the observed phenotypes. Meningeal- and subtype-specific drivers will be required to resolve these contributions. The source of regenerated meninges also remains unclear, and combining ablation with lineage tracing will help identify populations contributing to regeneration and how these responses change across development.

### Implications and future directions

Together, these findings establish the meninges as dynamic, heterogeneous tissues that coordinate brain and skull development across multiple stages in zebrafish. Given conservation of meningeal architecture and regulatory programs, this model may provide insight into human disorders involving meningeal-derived signals, including but not limited to craniosynostosis and calvarial dysplasia.

## Methods

### Zebrafish

All experiments were approved by the Institutional Animal Care and Use Committee at the University of California, Los Angeles (Protocol #ARC-2022-044). Published lines include Tg(*fli1a*:eGFP)^y1^ (Lawson and Weinstein, 2002), Tg(Hsa.*RUNX2*-Mmu.Fos:EGFP)^zf259^ (Kague et al., 2016), *Tg(−5.0kbfoxc1b:EGFP)^mw44^* (Miesfeld and Link, 2014), *Tg(foxc1b:gal4FF)^ca72^* (Whitesell et al., 2019)*, cdh1-mlanYFP^xt17^* (Cronan and Tobin, 2019)*, Tg(tbx18:DsRed2)^pd22^*(Kikuchi et al., 2011)*, Tg(UAS:GFP)(Asakawa et al., 2008), Tg(UAS-NTR:mCherry)^C264^* (Davison et al., 2007), *Tg(UAS:nlsGFP)^el609^* (Barske et al., 2020)*, osteoblast*:nlsEOS (Farmer et al., 2024), *foxc1a^el543^* (Xu et al., 2018)*, foxc1b^el620^* (Xu et al., 2018), Tg(sox10:CreERT)^el1777^ (Giovannone et al., 2019), and *Tg*(−3.5*ubb*:LOXP-EGFP-LOXP-mCherry) (Mosimann et al., 2011). Most experiments in this study were conducted prior to overt sex determination; however, when possible, equal numbers of males and females were used. One transgenic line was created using CRISPR/Cas9-based genomic integration of nlsEOS (Kimura et al., 2014; Thomas and Raible, 2019): *wu:fj16a03* (guide RNAs targeting the upstream of the 5’ UTR: 5′-AATGATGAAATATCTCTGCG-3). One-cell embryos were co-injected with guide RNAs (200 ng/μL), mbait guide RNA (200 ng/μL), IDT Cas9 protein (150 ng/μL), and a mbait-nlsEOS plasmid (20 ng/μL). Founders were identified by screening progeny for nlsEOS fluorescence. For tamoxifen treatments, embryos were collected at 1 dpf and treated overnight with 10 μM (Z)-4-hydroxytamoxifen (4-OHT; Sigma-Aldrich, H7904). Embryos were subsequently screened for mCherry expression at 5 dpf using a Zeiss fluorescence stereomicroscope.

### Single nuclei isolation

Adult zebrafish skullcaps were dissected away (*foxc1b*:gal4;UAS:nlsGFP, *fli1a*:GFP and DMSO and MTZ-treated *foxc1b*:Gal4;UAS:mCherryNTR) and fluorescent meninges attached to the brain were scraped away (*foxc1b*:gal4; UAS:nlsGFP) in Ringer’s solution. Tissue was mechanically minced with a razor and all tissue was transferred to a 1.5 mL tube for enzymatic dissociation (0.25% trypsin (Life Technologies, 15090-046), 1 mM EDTA, and 400 mg/mL Collagenase D (Sigma, 11088882001) in PBS). Tissue was incubated on a nutator at 28.5 °C for ∼45 min and the enzymatic reaction was stopped by adding 6X stop solution (6 mM CaCl2 and 30% fetal bovine serum (FBS) in PBS). Cells were pelleted by centrifugation (300 g) at 4 °C and resuspended in a final volume of 500 uL of 0.03% BSA in PBS. DAPI was added to resuspended cells, and cells were fluorescence-activated cell sorted to isolate live cells that were DAPI negative and GFP or mCherry positive for enrich for fibroblast populations or just DAPI negative cells for DMSO/MTZ skullcaps. Single nuclei suspension was collected by following 10X Genomics nuclei isolation protocol with low cell input for Multiome. FACS cells were pelleted for 15 min (300 rcf at 4°C) and incubated with lysis buffer for 90 s on ice. Isolated nuclei were washed by Wash buffer and Nuclei buffer and checked for nuclear integrity with a fluorescence confocal microscope with DAPI staining before use in library construction.

### Single cell-omic library construction, sequencing, and alignment

Multi-omic libraries (*foxc1b*:gal4;UAS:nlsGFP and *fli1a*:GFP) of snATACseq and snRNAseq from the same barcoded single nuclei were constructed per manufacturer’s instructions (10X Genomics, Chromium Next GEM Single Cell Multiome ATAC + Gene Expression). Single-Cell RNA sequencing libraries (DMSO or MTZ-treated *foxc1b*:Gal4;UAS:mCherryNTR) were constructed by 10X Genomics Chromium Single Cell 3′ Library and Gel Bead Kit v.2. To capture accessible chromatin and transcripts from the same nuclei in Multiome, accessible chromatin and polyadenylated mRNA from the same nuclei were pulled down and barcoded with the same sequences within isolated GEMs to achieve single-nuclei separation. cDNAs were synthesized through reverse transcription from polyadenylated mRNA. Quality control of libraries was assessed with the 4200 TapeStation system and Qubit dsDNA HS assay kit. Libraries were sequenced on Illumina NovaSeq (Multiome) and NextSeq (scRNAseq) platforms. For Multiome libraries, sequencing reads were aligned to customized genome built with GRCz11.fa, GRCz11.104.gtf, and JASPAR2022.pfm with addition of GFP gene information and called peak by Cell Ranger ARC v2.0.0 per manufacturer’s instructions to generate peak-by-cell and gene-by-cell count matrices. ScRNA-seq sequencing reads were aligned to customized genome built from GRCz11 with addition of mCherry gene information by Cell Ranger v3.0.0 to generate gene-by-cell count matrices.

### Single cell-omics data analysis

scRNA-seq data using the 10X Genomics Chromium Single Cell 3′ Library and Gel Bead Kit v.2 was analyzed as previously described (Farmer et al., 2024). Multi-omics data were analyzed by Seurat v5 and Signac v1.4 packages following the standard workflow with optimization. Cell Ranger ARC-output count matrices of individual libraries were used to create Seurat objects by “CreateSeuratObject” and “CreateChromatinAssay” functions. Quality control was performed by setting the thresholds of accessible region counts (nCount_ATAC) < 40000 (foxc1b) or < 30000 (fli1a), transcript counts (nCount_RNA) < 6000 (foxc1b) or < 7500 (fli1a), ratio of mononucleosome to nucleosome-free region (nucleosome_signal) < 3, and enrichment at TSS (TSS.enrichment) > 0.5. In order to compare accessible chromatin regions (peaks) between libraries, peaks-called were combined by “reduce” function that merges all intersecting peaks between libraries and filtered by new peak widths (peakwidths < 10000 & > 20). The reduced-peak list was used to recalculate peak-by-cell count matrices by “CreateFragmentObject”, “FeatureMatrix”, and “CreateChromatinAssay” functions. “Merge” function was used to combine count matrices of two libraries. Merged ATAC data were processed by “FindTopFeatures” (min.cutoff = 5) and LSI linear dimension reduction; and merged RNA data were processed by “SCTransform” and dimensionally reduced by PCA. 2:29 dimensions of LSI and 1:30 dimensions of PCA were used to cluster cells with both ATAC and RNA information by “FindMultiModalNeighbors” and “FindClusters” (resolution = 1, graph.name="wsnn", algorithm = 3). UMAP visualization was generated by “RunUMAP” (nn.name = "weighted.nn"). After filtering for artificial clusters of dying cells, 19 clusters were identified. Differentially enriched genes were calculated by “FindAllMarkers” (test.use = “wilcox”, logfc.threshold = 0.25, return.thresh = 0.01, min.pct = 0.25). Differentially accessible regions were calculated by “FindAllMarkers” (test.use = “LR”, logfc.threshold = 0.25, return.thresh = 0.01, min.pct = 0.1). To calculate motif enrichments of each cell by chromVAR, “AddMotifs” was first used to calculate motif enrichments of each accessible chromatin region (peak) followed by “RunChromVAR” to calculate motif activity in each cell. “FindAllMarkers” test.use = “wilcox”, mean.fxn = rowMeans, logfc.threshold = 0.25, return.thresh = 0.01, min.pct = 0.1) was performed to identify differentially enriched motifs. Enriched motifs from the same family were grouped together to calculate the frequency of enrichment for desired fibroblast clusters. Transcription factor motifs were obtained from the JASPAR2022 CORE vertebrate collection using the JASPAR2022 and TFBSTools R packages as previously described (Chen et al., 2023; Chen et al., 2025). Motif occurrences within accessible chromatin regions were identified using Signac, and chromVAR was used to calculate cell-level deviations in motif accessibility.

### Published dataset analysis

The processed Seurat object and associated annotation list were downloaded directly from the Daniocell website(Sur et al., 2023) and meninges clusters were subsetted and reclustered to capture transcriptionally distinct clusters by refining the resolution. For mouse datasets, individual leptomeningeal and dura mater datasets were downloaded from GEO and reanalyzed independently using standard Seurat methods. Module scores were calculated using the AddModuleScore function in Seurat with 10 control features. Scores were derived from 7–8 genes differentially expressed in zebrafish meningeal clusters with clear mouse homologs. FeaturePlots visualizing module scores were displayed with a minimum cutoff of the 10th percentile (q10) and a maximum cutoff of the 90th percentile (q90).

### RNAscope in situ hybridization and immunofluorescence

Embryonic and adult fish were measured and fixed individually in 4% PFA overnight at 4°C. Whole embryos processed in 10% sucrose in PBS for one day and 30% sucrose in PBS for at least one day before embedding tissue in Optimal Cutting Temperature (OCT) medium. Frozen sections were cut at 10 µm. Adult heads were decalcified for 2 weeks and processed by paraffin embedding as previously described (Farmer et al., 2024). RNAscope reagents were purchased from Advanced Cell Diagnostics, and experiments were performed using the RNAscope Fluorescent Multiplex V2 Assay according to the manufacturer’s protocol for formalin-fixed paraffin-embedded (adult) or for fixed frozen (embryos) sections. For cryosections, heat antigen retrieval was omitted. The following probes were used for this study: Channel 1, dr-*gttcb*, dr-*krt94*, dr-*slc6a2,* dr-*foxc1b, dr-tbx18*, *dr-stra6*; Channel 2, dr-*coch*, dr-*foxc1a*, *mcherry*, dr-*slc4a7*; Channel 3, dr-*ccn2a*, dr-*igfbp2a*; Channel 4, dr-*rspo2*. RNAscope experiments were followed by immunofluorescence or performed independently using the same antigen retrieval reagents as the RNAscope Fluorescent Multiplex kit, and slides were blocked for 1 h in 10% (adult stains) or 5% (embryonic stains) goat serum, stained with primary overnight (anti-laminin, 1:250, Millipore #L9393; anti-GFP, 1:1000, Novus NB100-1614; anti-mCherry, 1:1000, Novus NBP2-25157) and stained with Alexa-Fluor secondaries from Thermo Scientific (1:250, Thermo) and Hoechst 33342 at room temperature for 1 h. Wholemount staining of adult brains were performed by permeabilizing tissues with 0.25% Triton for 3 hours at room temperature followed by blocking in anti-acetylated α-tubulin antibody (1:500, Sigma T6793) overnight, and the secondary for two hours at room temperature with secondary antibody and Hoechst 33342.

### Plastic embedding

Zebrafish heads were dissected and fixed in 70% ethanol without decalcification. After dehydration, samples were gradually infiltrated with increasing concentrations of methyl methacrylate resin and embedded in 2.5% methyl methacrylate. Longitudinal sections (5 µm thick) were obtained using a Polycut microtome fitted with a tungsten carbide knife and stained with toluidine blue for histological analysis. Sections were subsequently stained with Toluidine Blue staining and imaged using an ECHO Revolution microscope.

### 2-photon ablation

5 dpf larvae were mounted in low-melting-point agarose and imaged on an inverted LSM 880 microscope equipped with a two-photon laser. Using a 20× air objective, a region at the midbrain– hindbrain boundary was manually delineated using the ROI tool to encompass approximately 20 nuclei. Targeted ablation was performed at 450 mW with a scan speed of 400 Hz for a total duration of 9 s (three cycles of 3 s each). Ablation efficacy was assessed by imaging in the transmitted photomultiplier tube (T-PMT) channel to confirm loss of pigmented cells. Z-stacks were acquired immediately after ablation and again 6 h post-treatment to verify sustained signal loss. Following imaging, larvae were individually housed and re-imaged at 1 and 2 days post-treatment (dpt) before being raised to adulthood.

### Adult skeletal preparations

Skeletal preparations were performed as previously described (Teng et al., 2018), and zebrafish were imaged on a Zeiss Stemi with Zeiss Labscope software in 100% glycerol.

### Imaging

All imaging was performed on a Zeiss LSM880 or a Zeiss LSM980 microscope using ZEN software. For repeated live imaging experiments, fish were anesthetized in Tricaine, mounted, and imaged using a 10X objective. For whole calvaria nlsEOS conversion experiments, live fish were transferred to a 6- or 12-well dish and exposed to a handheld UV light (UV Flashlight Black Light, 3-in-1 Magnetic Flashlight Rechargeable, AdamStar) for 5–30 min. Converted fish were individually housed and re-imaged for up to 2 months following initial conversion.

### Metronidazole treatment

Zebrafish carrying *foxc1b*:gal4;UAS:mCherryNTR were outcrossed to wildtype AB zebrafish and sorted by 5 dpf. mCherry^+^ fish were split into two groups to receive DMSO in Embryonic Media (EM) or 10 mM Metronidazole in EM. Larvae were treated overnight and washed 3 times in EM before entering the nursery at 6 dpf. Tanks for both conditions were assessed daily to assess survival rate and imaged to confirm loss or return of fluorescence signal.

## Inhibitor drug screen

Zebrafish embryos were screened for *foxc1b:*Gal4;UAS:mCherryNTR fluorescence at approximately 18 hpf and chorions were digested enzymatically with 2 mg/mL Pronase. Fish carrying *tbx18:*DsRed2 could not be screened before drug treatments. DAPT (Tocris, 2634), 4-Diethylaminobenzaldehyde (Sigma-Aldrich, D86256), Dorsomorphin (Selleckchem S7840), Infigratinib (Selleckchem, S2183), LY2109761 (Selleckchem, S2704), XAV-939 (Selleckchem, S1180), and IWR-1-Endo (Selleckchem, S7086) were dissolved in DMSO and aliquoted as stock solutions. DAPT, Infigratinib, and LY2109761 stock solutions were added to heated E3 and cooled to room temperature before treating embryos. DEAB and Dorsomorphin stocks were diluted in room temperature Embryo Medium. XAV-939 and IWR-1-Endo stocks was added to heated Embryo Medium and cooled to room temperature. Cyclopamine (Selleckchem S1146) was dissolved in Ethanol, and stock was diluted into heated E3 and cooled to room temperature. PTU media was added to all drug solutions, regardless of whether prepared in EM or E3, to a final concentration of 197 μM. Fresh drug was added daily for animals treated from 24-72 hpf. For transient inhibition experiments, drug media was replaced with 197 μM PTU in embryo medium every 24 hours until experiment termination. Animals were euthanized and fixed in 4% PFA at 4°C overnight prior to imaging.

### Quantification and statistical analysis

For *foxc1a/*b allelic series, regions were defined by manually drawing ROIs around brain domains. Fluorescent signal in the brain or deeper tissues was removed from the z-stack to avoid misidentifying as dorsal meninges signal. Max projections of edited z-stacks were then thresholded and measured for fraction of mCherry^+^ pixels using ImageJ. For quantifying mCherry expression around the perimeter of 5 dpf cryosections, brain domains and mCherry^+^ meninges were manually defined in ImageJ by establishing ROI perimeters in ImageJ. For meninges ablation experiments, entire calvaria were defined by manually drawing a ROI around the frontal and parietal bones using ImageJ. Hole regions were defined by manually drawing a ROI around the area where bone was absent in the calvaria. The measure tool was then used to obtain the areas of the entire calvaria and the hole regions in pixels. To quantify the size of each hole relative to the whole calvaria size, a ratio of hole area divided by entire calvaria area was calculated for each sample. For osteoblast:nlsEOS experiments, regions were defined by manually drawing a ROI across bone fronts or cranial sutures using ImageJ. The channels were then split and nuclei within each channel were counted manually. De novo osteoblasts were defined as green only cells. Colocalization analysis was performed in Fiji/ImageJ using the Coloc 2 plugin. Regions of interest encompassing the relevant signal were selected from matched fluorescence channels, and colocalization was quantified using thresholded Manders coefficients (tM1 and tM2). tM1 represents the fraction of signal in channel 1 overlapping thresholded signal in channel 2, whereas tM2 represents the reciprocal fraction of channel 2 overlapping channel 1. Thresholds were determined within Coloc 2, and coefficients were reported without background subtraction. Non-overlapping regions were analyzed as negative controls to estimate baseline colocalization. For each comparison, tM1 and tM2 were reported separately because the two coefficients reflect distinct directional measurements of signal overlap. All statistical analyses were completed using PRISM.

## Data availability

The raw and processed sequencing data for single-cell genomics datasets in this study are available on GEO (accession number: GSE318179).

## Acknowledgements

The authors thank the Bone Histomorphometric Laboratory and Broad Stem Cell Research Center’s Flow Cytometry Core and Sequencing Core for their services. We thank you Susan Childs for providing the crucial *foxc1b*:Gal4;UAS:mCherryNTR fish line.

## Contributions

A.L.A, K.W., H.C., and D.T.F conceived and designed the study. A.L.A. and B.M.H. performed larval experiments. K.W. and D.T.F. performed adult experiments. P.R. and K.W. generated and described the *wu:fj16a03*:nlsEOS reporter. H.C performed single-cell and bioinformatic analysis. D.T.F. supervised the research. D.T.F. wrote the manuscript with support from A.L.A, K.W., and H.C.

