## Supplemental Information for "Conserved cellular architecture and developmental mechanisms of the zebrafish meninges"

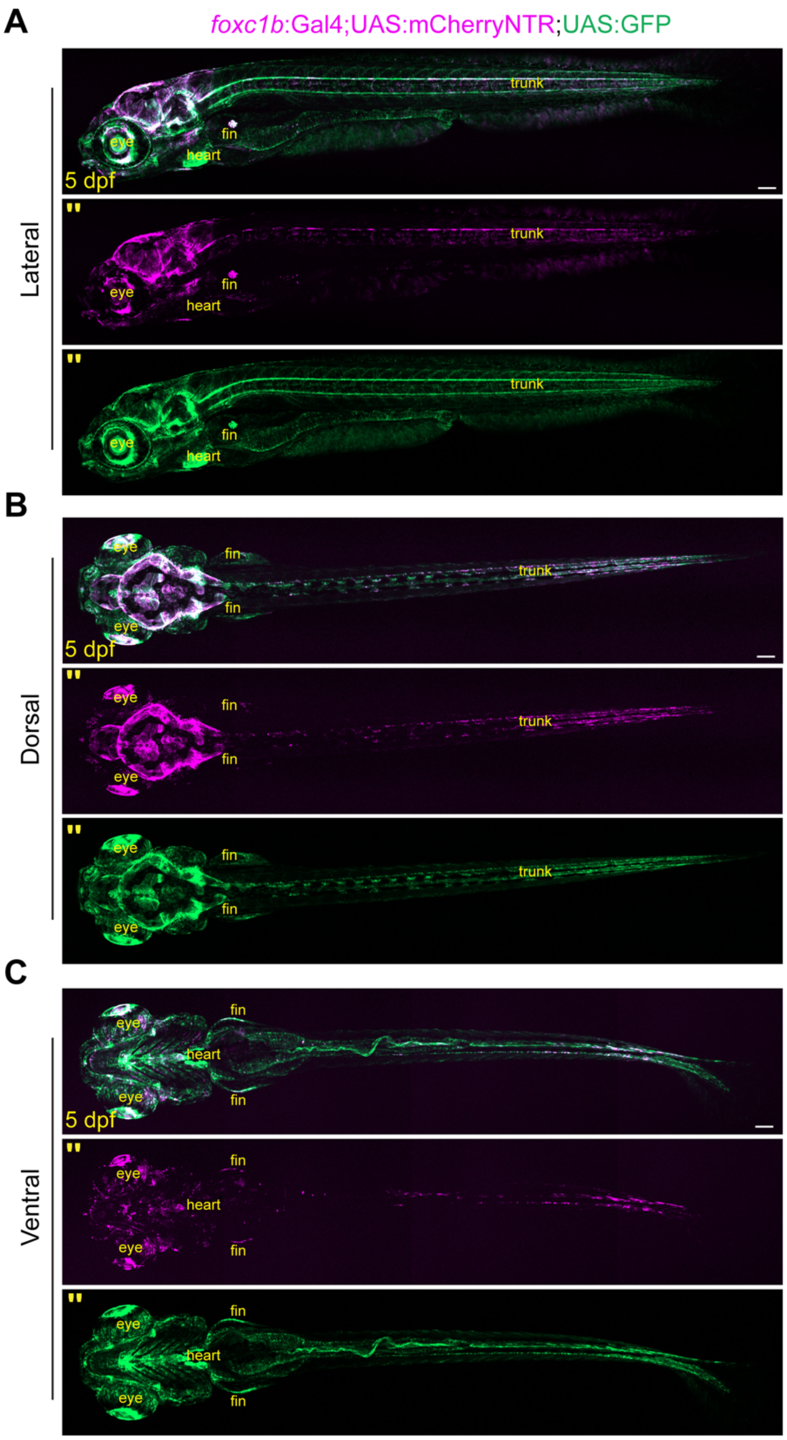


**Figure S1. Whole-body imaging of the *foxc1b*:Gal4 reporter. (A-C)** Lateral, dorsal and ventral view confocal images of larvae carrying foxc1b:Gal4;UAS:mCherryNTR with UAS:GFP. (") = single channels. Scale bars, 100 µm.

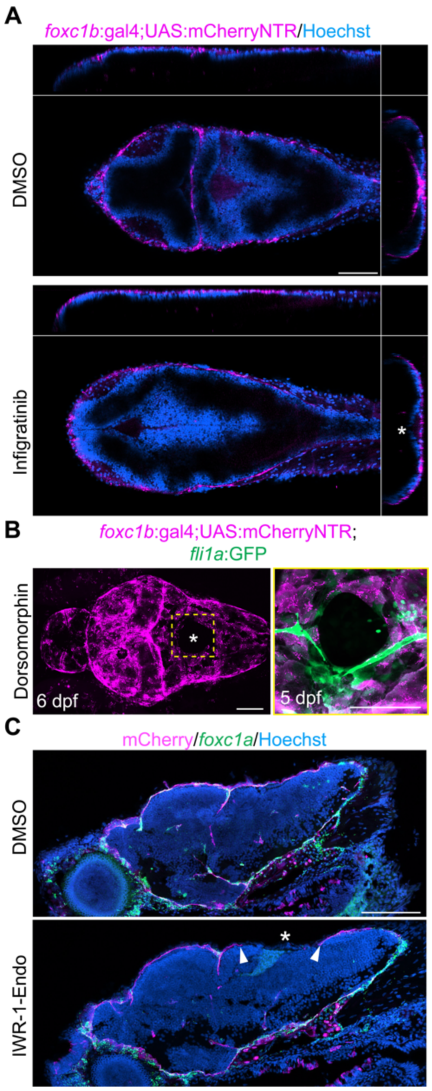


**Figure S2. Impaired anatomy and meninges establishment after pathway inhibition. (A)** Confocal images of DMSO- and Infigratinib-treated larvae carrying *foxc1b*:Gal4;UAS:mCherryNTR and stained with Hoechst , with orthogonal views shown above (XZ) and to the right (YZ). Asterisk indicates reduced mCherry$^{+}$ signal between the midbrain and hindbrain (n = 3 per condition). **(B)** Dorsal-view confocal images of *foxc1b*:Gal4;UAS:mCherryNTR larvae treated with Dorsomorphin using the transient treatment paradigm (Fig. 3E) and imaged at 5 or 6 dpf (n = 10 larvae per condition). Asterisk marks rostral regions where mCherry$^{+}$ signal is reduced. The boxed region is a higher magnification image that includes co-imaging with *fli1a*:GFP. **(C)** RNAscope analysis of *foxc1a* combined with mCherry immunostaining in *foxc1b*:Gal4;UAS:mCherryNTR larvae treated with DMSO or IWR1-Endo (n = 3 per condition). Arrowheads mark *foxc1a*$^{+}$mCherry$^{+}$ regions, and asterisks indicate reduced mCherry$^{+}$ signal. Scale bars, 100 μm.

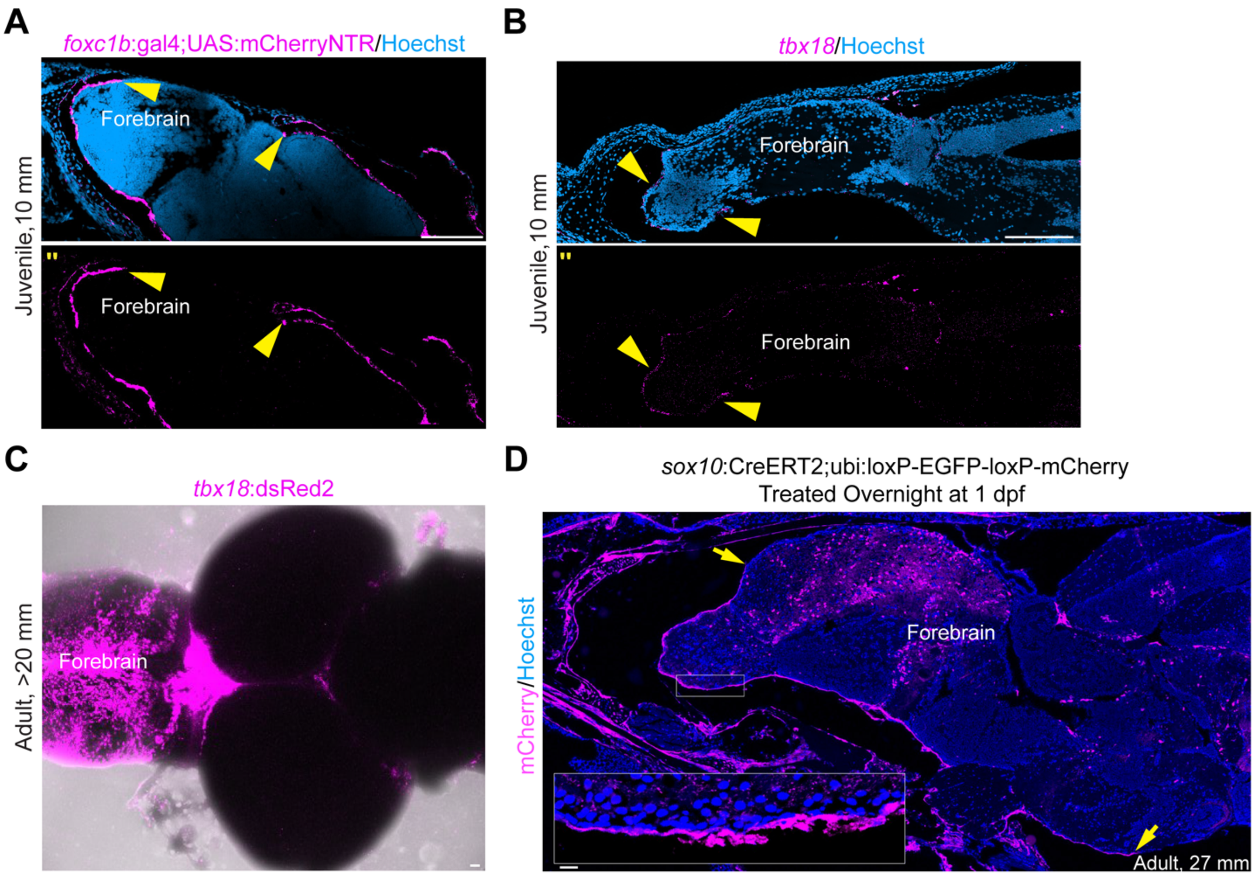


**Figure S3. Evaluation of forebrain meninges at juvenile and adult stages. (A)** Immunostaining for mCherry in a *foxc1b*:Gal4;UAS:mCherryNTR juvenile at the medial dorsal forebrain meninges (n = 3). Arrowheads mark the boundaries of mCherry$^+$ signal. **(B)** RNAscope analysis of *tbx18* in the forebrain (n = 2). Arrows mark *tbx18*$^+$ signal around the forebrain. **(C)** Dorsal view of a dissected brain from adult *tbx18:*dsRed2 fish, with brightfield imaging shown for context (n = 4). **(D)** Immunostaining for mCherry on sections from a *sox10*:CreERT2;ubi:loxP-EGFP-loxP-mCherry animal treated 1 dpf (n = 2). Arrowheads indicate mCherry$^+$ cells around the forebrain. Boxed region shows a higher-magnification view of the representative area. (") = single channels. Scale bars, 100 µm.

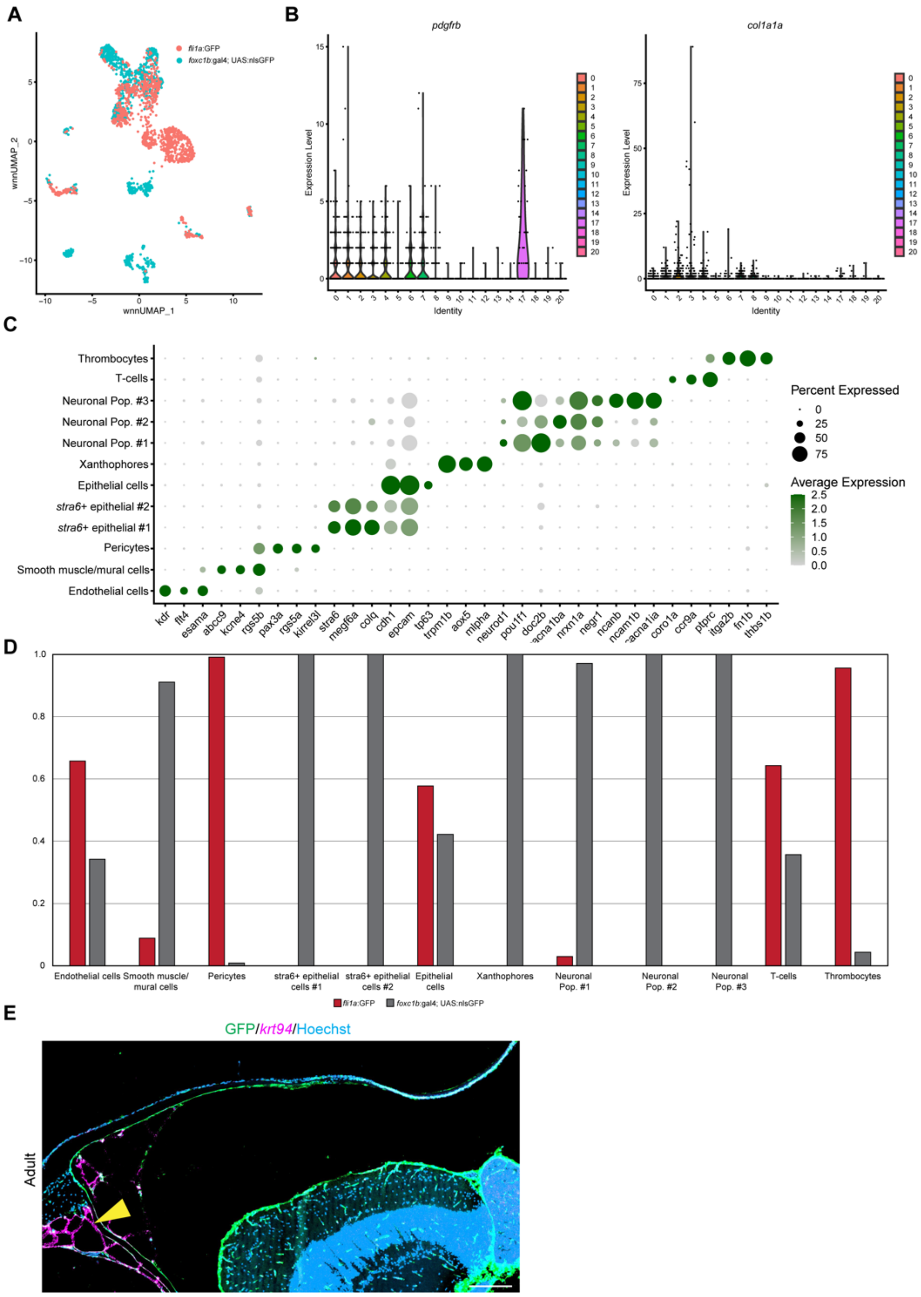


**Figure S4. Analysis of non-meningeal fibroblast clusters. (A)** Visualization of merged single-cell datasets colored by dataset of origin. **(B)** Violin plots of *pdgfrb* and *col1a1a*. **(C)** Dot plot showing marker genes enriched in each non-fibroblast clusters within the merged dataset. **(D)** Graphical depiction of the fraction of each non-fibroblast cluster captured from each dataset. **(E)** RNAscope analysis for *krt94* combined with GFP immunostaining in adult *foxc1b*:Gal4;UAS:GFP animals. Arrowhead indicates GFP$^{+}$*krt94*$^{+}$ cells. Scale bars, 100 µm.

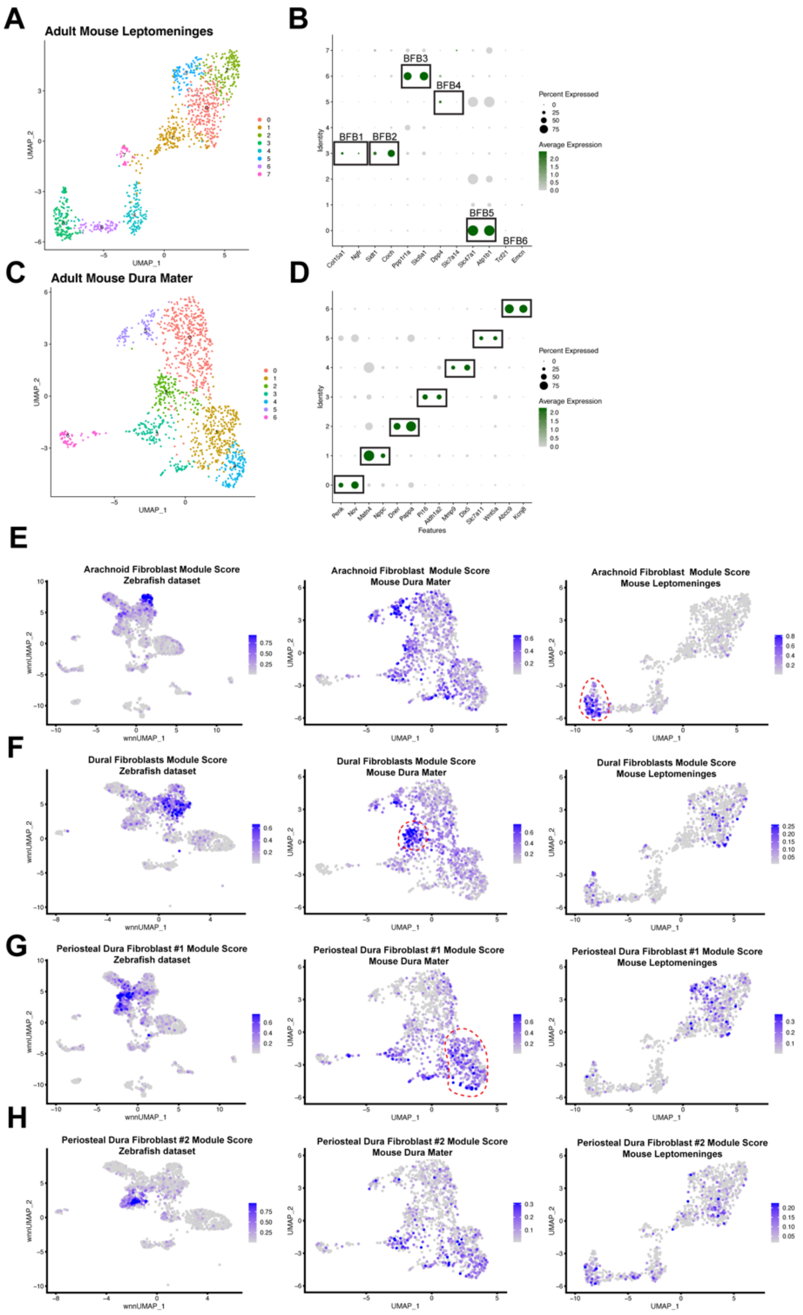


**Figure S5. Mapping gene modules reveals shared transcriptional profiles between zebrafish and mouse meninges. (A)** UMAP clustering of the leptomeninges single-cell dataset from Pietilä et al., 2023. **(B)** Dot plot showing marker gene expression across cell types defined in Pietilä et al., 2023. **(C)** UMAP clustering of the dura mater single-cell dataset from Pietilä et al., 2023. **(D)** Dot plot showing marker genes enriched in each cluster identified in (C). **(E–H)** Module scores calculated from genes enriched in each zebrafish meningeal cluster and mapped to the zebrafish dataset, with corresponding mouse paralogs mapped to both the leptomeninges and dura mater datasets.

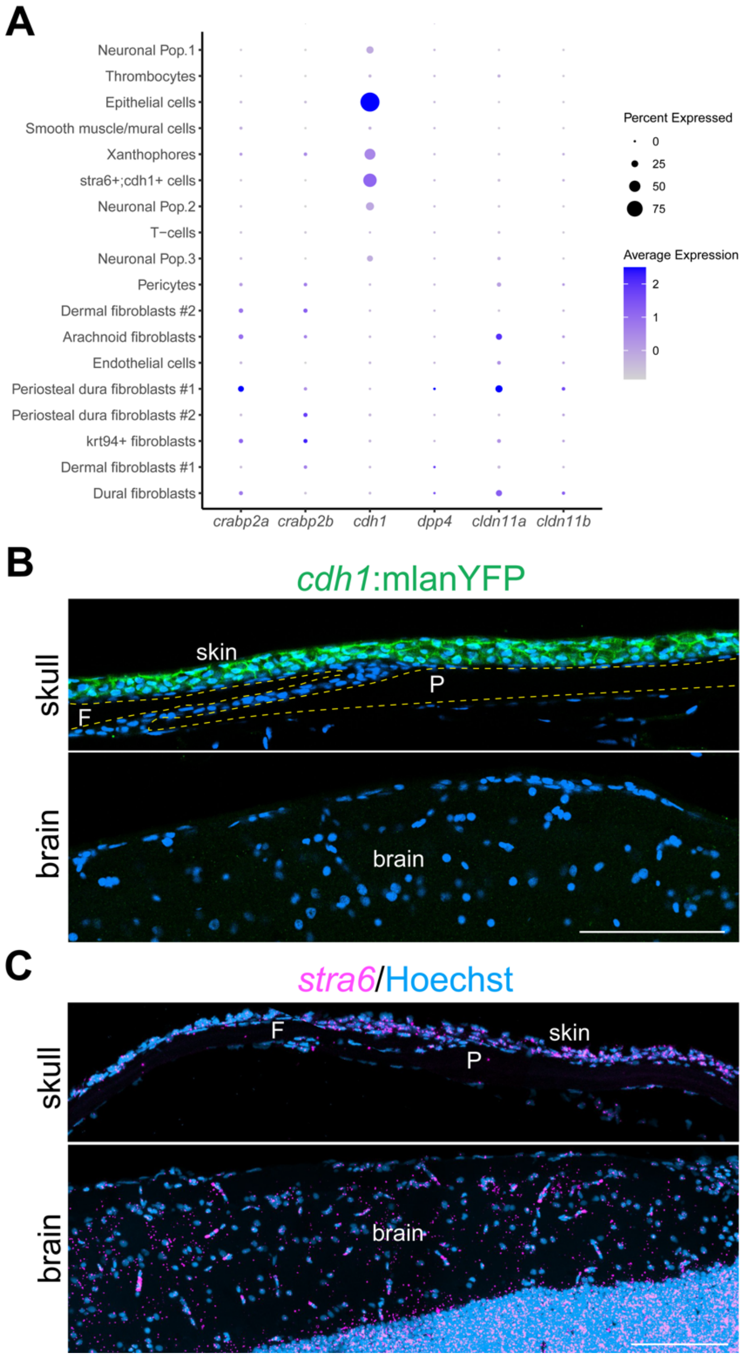


**Figure S6. Markers for arachnoid barrier cells are not specifically enriched within zebrafish meninges datasets. (A)** Dot plot showing expression of marker genes for arachnoid barrier cells defined in Pietilä et al., 2023. **(B)** Immunostaining for YFP using a GFP antibody in *cdh1*:mlanYFP zebrafish. **(C)** RNAscope analysis of *stra6* in the skullcap and brain. Scale bars, 100 μm.

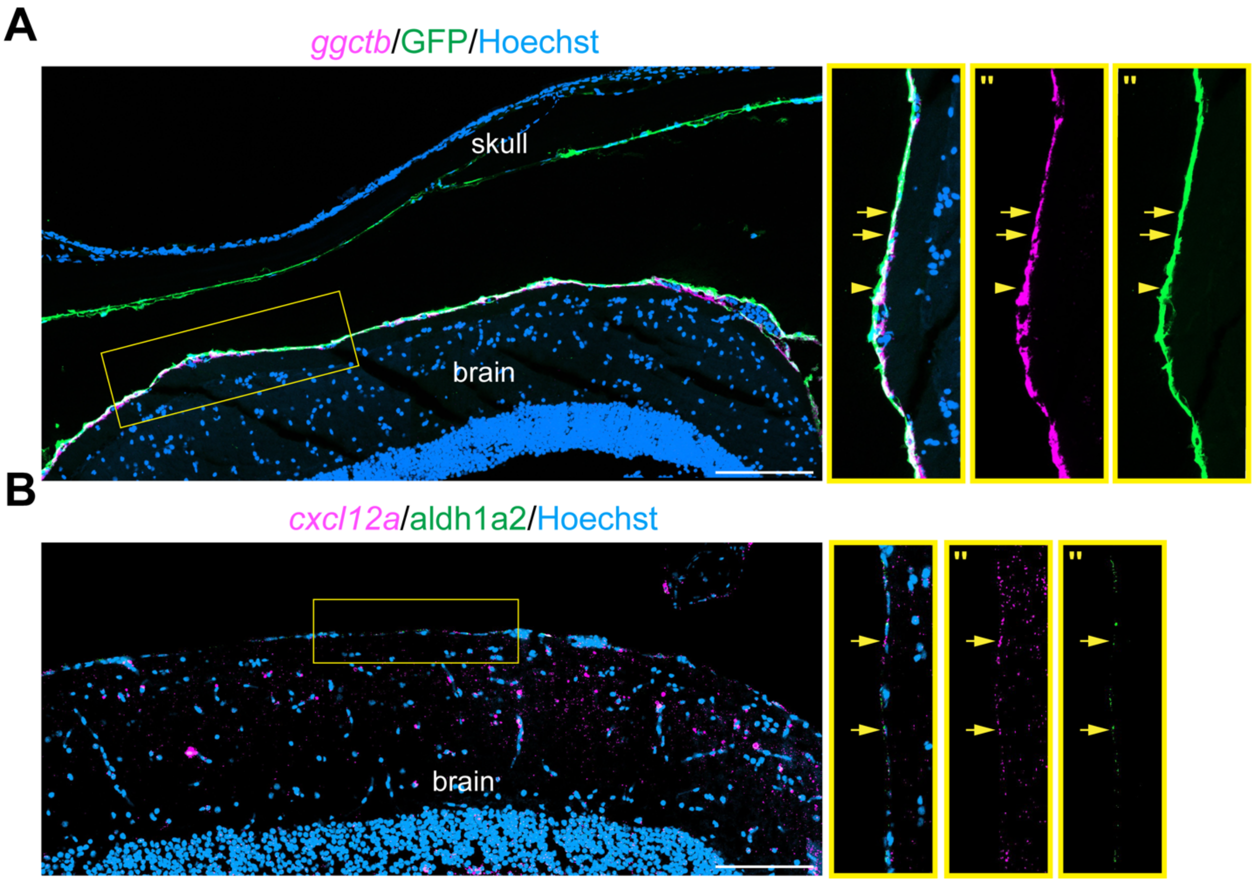


**Figure S7. Expression of the leptomeningeal marker *ggctb* and pial associated genes within *foxc1b* reporter-positive cells. (A)** RNAscope analysis for *ggctb* combined with GFP immunostaining in adult *foxc1b*:Gal4;UAS:GFP animals shows localized detection of *ggctb*-positive, GFP-positive cells (arrows) surrounding the brain with scattered GFP single positive cells (arrowhead). **(B)** RNAscope analysis for *cxcl12a* and *aldh1a2* reveals expression (arrows) in meninges adjacent to the brain. Boxes indicates magnified region corresponding to the location highlighted in the image. All stains were performed in biological triplicates. (") = single channels. Scale bars, 100 μm.

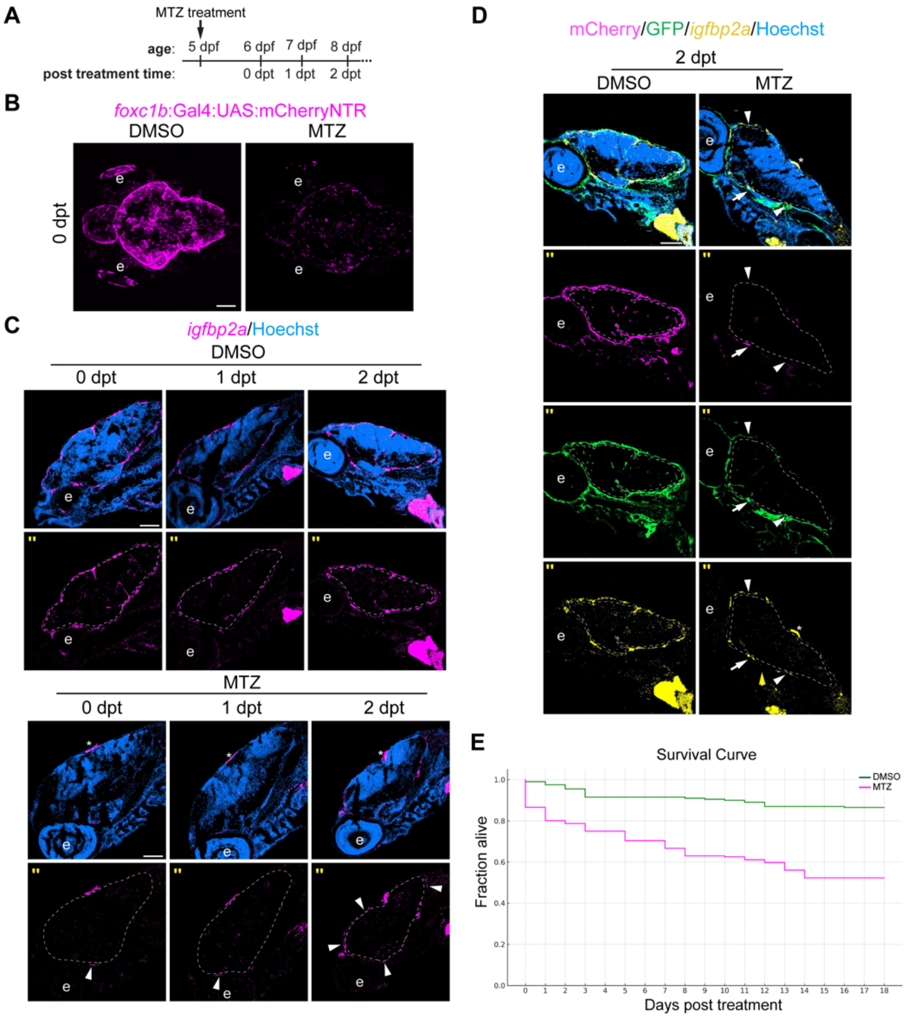


**Figure S8. Larval zebrafish survive widespread meningeal ablation and rapidly regenerate.** **(A)** Schematic illustrating the drug treatment strategy. **(B)** Dorsal-view confocal images of *foxc1b*:Gal4;UAS:mCherryNTR embryos treated with DMSO or MTZ (n = 5 larvae per condition). **(C)** RNAscope analysis for *igfbp2a* in *foxc1b*:Gal4;UAS:mCherryNTR larvae at 0, 1, or 2 days post-treatment (dpt) shows gradual re-emergence of *igfbp2a*-positive cells following MTZ treatment (n = 3 larvae per condition). The yellow dashed line outlines the brain. Arrowheads indicate *igfbp2a*-positive domains in treated larvae, and asterisks mark *igfbp2a* expression domains not associated with the meninges. **(D)** RNAscope analysis of *igfbp2a* combined with mCherry and GFP immunostaining in adult *foxc1b*:Gal4; UAS:mCherryNTR; UAS:GFP animals treated with DMSO or MTZ (n = 3 larvae per condition). The yellow dashed line outlines the brain. Arrowheads indicate $GFP^{+}$*igfbp2a*$^{+}$ cells and arrow indicates $mCherry^{+}GFP^{+}$*igfbp2a*$^{+}$ cells. **(E)** Survival curve for DMSO- and MTZ-treated zebrafish. e = eye. ('') = single channels. Scale bars, 100 µm.

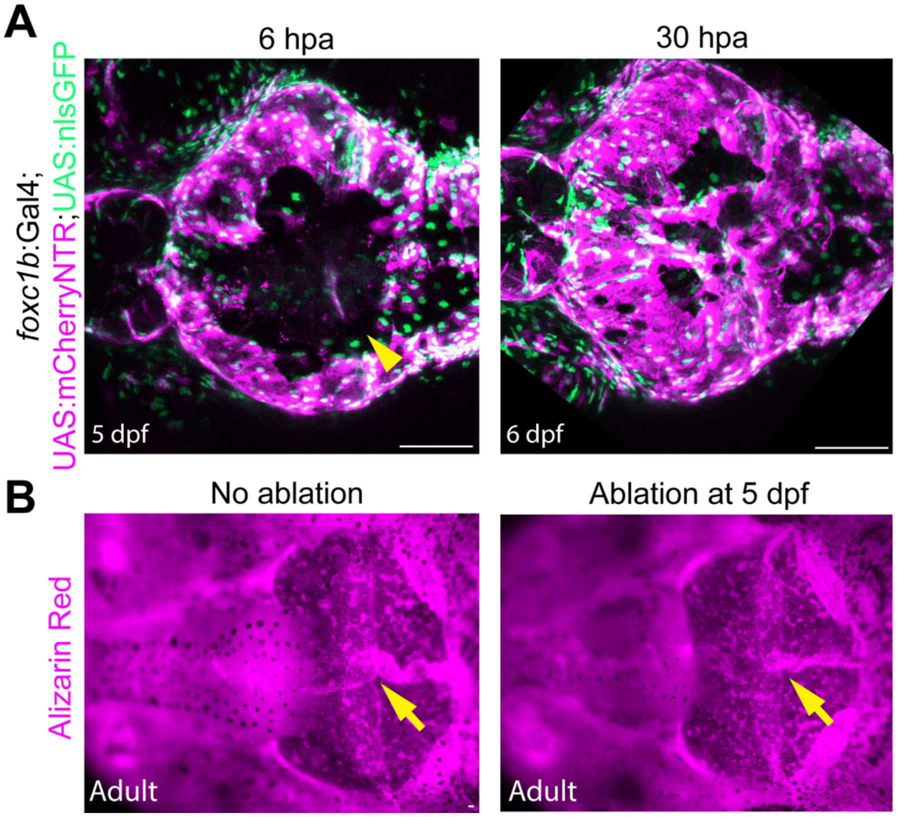


**Figure S9. Local ablation of larval meninges does not impair meningeal or calvarial development. (A)** Dorsal-view confocal images of *foxc1b*:Gal4;UAS:mCherryNTR;UAS:nlsGFP larvae at 6 hours post-ablation (hpa) and 30 hpa show transient loss of meningeal cells followed by rapid regeneration (n = 8 fish per condition). Arrowhead marks ablated region with no fluorescent signal. **(B)** Dorsal views of Alizarin-stained skulls from control and ablation-treated adult zebrafish reveal no detectable changes in calvarial morphology (arrows) (n = 5 fish per condition). Scale bars, 100 µm.

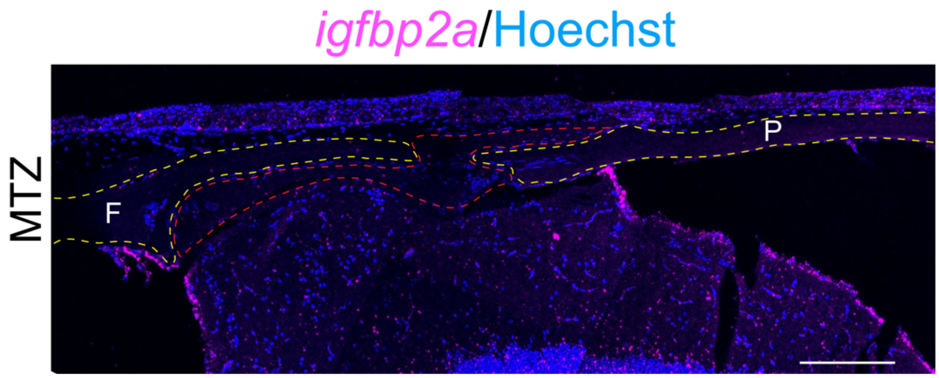


**Figure S10. Meningeal generation is impaired at sites of ectopic bone formation.** RNAscope analysis for *igfbp2a* in MTZ-treated animals with ectopic bone formation shows that bone develops over regions lacking detectable *igfbp2a* expression within the skull. Yellow dashed lines mark the presumptive frontal and parietal bones. Red dashed line outlines the ectopic bone. All stains were performed in biological triplicates. F= frontal bone, P= parietal bone. Scale bars, 100 µm.

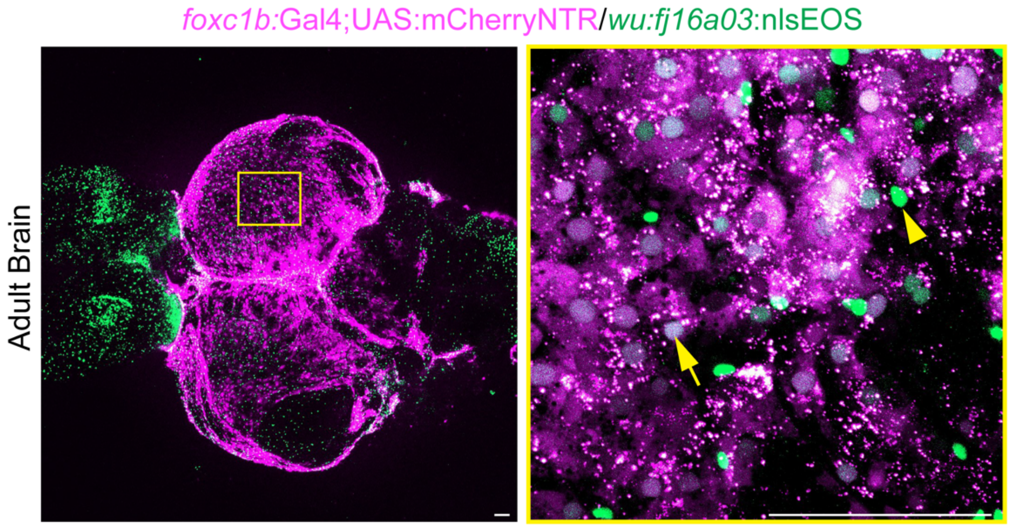


**Figure S11. Overlap between *wu:fj16a03*:nlsEOS and the *foxc1b* reporter in adult meninges.** Dorsal-view confocal images of dissected adult brains show partial co-detection of *foxc1b*:Gal4;UAS:mCherryNTR and *wu:fj16a03*:nlsEOS signals (n = 5). Higher-magnification (right) view highlights double-positive cells (arrow) as well as *wu:fj16a03*:nlsEOS single-positive cells (arrowhead). Scale bars, 100 μm.

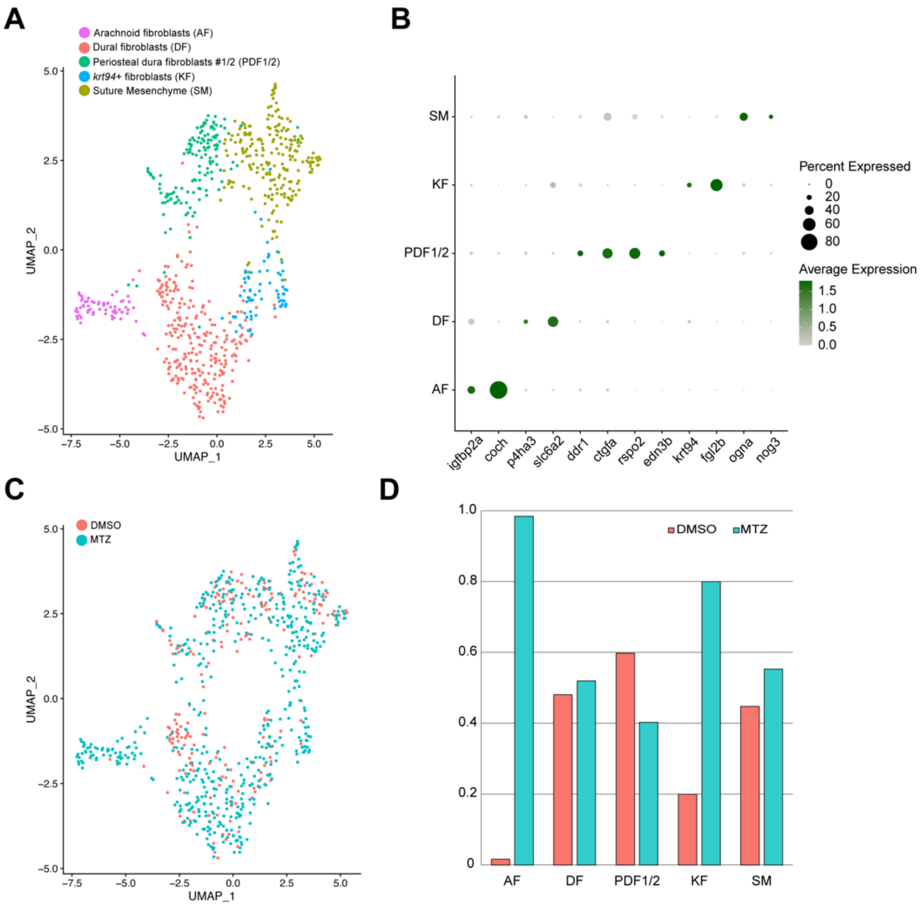


**Figure S12. Single-cell RNA sequencing identifies brain-associated meningeal population attached to MTZ-treated skullcaps. (A)** UMAP clustering of fibroblast cells isolated from DMSO- and MTZ-treated skullcaps. **(B)** Dot plot showing marker genes enriched in each cluster identified in (A). **(C)** Visualization of merged single-cell datasets colored by dataset of origin. **(D)** Graphical depiction of the fraction of each cluster captured from each dataset.

**Table S1.** Co-localization analysis for indicated figures

**Table S2.** Differential gene expression of merged adult dataset from Multiome analysis

**Table S3.** Differential gene expression of merged adult skullcaps from DMSO and MTZ scRNA-seq

**Movie 1.** Time-lapse imaging of an embryo carrying *fli1a*:GFP and *foxc1b*:Gal4;UAS:mCherryNTR reporters, beginning at 24 hpf and continuing for approximately 72 hours. Development may be delayed due to prolonged imaging.

**Movie 2.** Time-lapse imaging of a *foxc1a*$^{-/-}$; *foxc1b*$^{-/-}$ embryo carrying *fli1a*:GFP and *foxc1b*:Gal4;UAS:mCherryNTR reporters, beginning at 24 hpf and continuing for approximately 72 hours. Development may be delayed due to prolonged imaging.
